# Selective Ablation of 3’ RNA ends and Processive RTs Facilitate Direct cDNA Sequencing of Full-length Host Cell and Viral Transcripts

**DOI:** 10.1101/2022.01.27.478099

**Authors:** Christian M. Gallardo, Anh-Viet T. Nguyen, Andrew L. Routh, Bruce E. Torbett

## Abstract

Alternative splicing (AS) is necessary for viral proliferation in host cells and a critical regulatory component of viral gene expression. Conventional RNA-seq approaches provide incomplete coverage of AS due to their short read-lengths and are susceptible to biases and artifacts introduced in prevailing library preparation methodologies. Moreover, viral splicing studies are often conducted separately from host cell transcriptome analysis, precluding an assessment of the viral manipulation of host splicing machinery. To address current limitations, we developed a quantitative full-length direct cDNA sequencing strategy to simultaneously profile viral and host cell transcripts. This nanopore-based approach couples processive reverse transcriptases with a novel one-step chemical ablation of 3’ RNA ends (termed CASPR) which decreases ribosomal RNA reads and enriches for poly-adenylated coding sequences. We extensively validate our approach using synthetic reference transcripts and show CASPR doubles the breadth of coverage per transcript and increases detection of long transcripts (>4kb), while being functionally equivalent to PolyA+ selection for transcript quantification. We used our approach to interrogate host cell and HIV-1 transcript dynamics during viral reactivation and identified novel putative HIV-1 host factors containing exon skipping or novel intron retentions and delineated the HIV-1 transcriptional state associated with these differentially regulated host factors.

## Introduction

Alternative splicing (AS) greatly increases protein diversity encoded by the human genome, and has been estimated to occur in up to 95% of genes with multiexonic transcripts (1). This process is tightly regulated by cis- and trans-acting elements, chromatin accessibility, and other signaling pathways (2). Alternative splicing has been shown to be a driver of human proteome diversity (3,4) and a critical regulatory component in the tissue-specific expression of human transcriptomes(5). Recently, increasing use of massively parallel RNA-Seq pipelines have allowed population-scale transcriptome studies which have revealed naturally occurring variants that modulate AS and influence disease susceptibility(6).

Viral infections commonly alter host cell splicing landscape, as shown by genes that appear differentially-spliced upon viral infection in transcriptomic studies, or splicing-related genes that appear differentially enriched or phosphorylated in proteomic studies (7). In cells infected with HIV-1 (HIV), alternatively-spliced host cell transcripts have been shown to promote a permissive environment for viral activation and proliferation via induction of alternative transcription start/end sites (8) and via functional enrichment of HIV replication related pathways (9). Similarly, proteomic studies have shown induction of signaling pathways involved in mRNA splicing in T-lymphocytes upon HIV entry (10), with phosphorylation of canonical splice factors being the apparent regulatory mechanism. Additionally, splicing-related host factors have been reported which bind HIV accessory proteins and act as trans-regulatory elements including the binding of U2AF65 and SPF45 by Rev (11) and SR proteins by Vpr (12), as well as the interactions between POLR2A and Tat (13).

Alternative splicing is also a critical regulatory mechanism of viral gene expression (14). In HIV, a single unspliced 9.2kb RNA serves as both the genome and mRNA for both Gag and Gag-Pol polyproteins, while alternatively-spliced mRNA variants code for the 7 remaining gene products by dynamically and specifically interacting with regulatory elements, thereby generating over 50 physiologically relevant transcripts that can be grouped in ‘partially spliced’ (4kb) and ‘completely spliced’ (1.8kb) groups (15-18). The underlying mechanism in AS regulation of HIV transcripts is the placement of the open-reading frames of each gene in close proximity to the single transcription start site region at the 5’ end of HIV-RNA, thus optimizing the coding potential of HIV-genes by translating different proteins from a common mRNA. The completely spliced 1.8kb class is particularly important during the early infection phase, and it includes Tat and Rev transcripts which respectively aid in transcription and export of partially spliced transcripts from the nucleus. An eventual shift in splicing dynamics, partially attributed to Rev, results in increased production of partially spliced and unspliced mRNAs (11). Thus, carefully orchestrated splicing dynamics are critical for regulating the dynamics of HIV gene expression and any resulting interactions with host factors.

Conventional RNA-seq approaches, while robust and reproducible, are limited by their read-length in providing full coverage of AS events (such as alternate donor/acceptor sites, exon skipping, alternate exon usage, and intron retention). Moreover, library preparation techniques can introduce biases/artifacts due to PCR amplification bias, artefactual recombination, fragmentation, or targeted enrichment methods for coding sequences (CDS) (19). The read-length limitation in short-read RNA-seq, coupled with the biases and artifacts introduced in prevailing library preparation methodologies can prevent an assessment of full exon connectivity in a quantitative manner, resulting in loss of information on transcript isoform diversity, including splice variants (20). The limitations of current RNA-seq approaches are particularly exacerbated when assessing transcript expression in polycistronic HIV RNA where all transcripts are flanked by identical 5’ and 3’ end exons (only varying in their internal splicing sites) and vary greatly in overall transcript length. Previous attempts to address these constraints have used primer sets for each transcript class or gene product, relied on molecular barcoding, or used emulsion PCR to ameliorate PCR skewing or sampling biases (18,21). However, use of different primer sets prevents the quantitative comparison between transcripts and does not provide full exon coverage, while molecular barcoding approaches were used with short-read NGS approaches. Previous HIV splicing studies were not implemented within the context of a host cell transcriptome analysis, precluding a direct assessment of the viral manipulation of host splicing machinery or further insights into virus-host interaction dynamics (22). Since the regulation of HIV gene expression depends on the ability of the virus to co-opt host cell splicing machinery, understanding host cell transcriptional state and its resulting HIV mRNA-splicing signature would identify novel molecular signatures of HIV infection and provide opportunities for drug/probe development based on novel viral/host factor interactions.

To address current RNA-seq limitations, we developed and validated a quantitative full-length RNA-seq strategy for the simultaneous profiling of poly-adenylated host and viral transcripts from unamplified cDNA. This nanopore sequencing based approach is supported by use of processive reverse transcriptases (RTs), and oligo-d(T) priming, coupled with a one-step chemical ablation of 3’ RNA ends (named CASPR) which decreases ribosomal RNA reads and enriches for poly-adenylated transcripts. We validate both RT conditions and CDS enrichment strategies using synthetic reference transcripts and show that while CASPR is functionally equivalent to PolyA+ selection for transcript quantification purposes, it provides critical advantages in doubling the breadth of coverage per transcript and significantly increasing the efficiency of capture of long transcripts >4kb in size. This improved practical throughput and likelihood of capturing full-exon connectivity. We then demonstrate the utility of our pipeline by interrogating host cell and HIV transcript dynamics in reactivated J-Lat 10.6 cells, a widely-used cell-line model of HIV reactivation (23,24). We identify putative host factor correlates of HIV transcriptional reactivation that contain exon skipping events (PSAT1) or novel intron retentions (PSD4) and delineate the HIV transcriptional state associated with these differentially regulated host factors. We anticipate this pipeline will allow greater insights into host cell-pathogen transcript dynamics involved in viral infection and activation.

## Materials and Methods

### Cell Culture

J-Lat 10.6 cells, a Jurkat-derived cell line that is latently infected with HIV(23), were obtained from the NIH AIDS Reagent Program (clone #10.6, Dr. Eric Verdin). The J-Lat 10.6 clone contains a single R7/ΔEnv strain integrated into the SEC16A locus, and EGFP inserted into the nef ORF. For control experiments, the Jurkat E6-1 clone was obtained from the NIH AIDS Reagent Program (cat #177, from Dr. Arthur Weiss(25)). J-Lat 10.6 cells were activated with 10ng/mL TNF-alpha (PeproTech 300-01A) for 24 hours which induces latency reversal of integrated provirus, resulting in positive GFP expression and p24 production which are respectively detected via flow cytometry and p24 ELISA. Cell lines were maintained in RPMI 1640 (Life Tech) supplemented with 10% FBS (Hyclone) and 1% Pen/Strep at 37°C and 5% CO_2_.

### Total RNA isolation

Total RNA was isolated from cell pellets (<1×10^7^ cells) using the RNeasy Mini kit (QIAGEN, cat. 74134). Cells were lysed with RLT buffer (with no ß-ME) and processed according to manufacturer’s instructions, and eluted in 25-50 µL nuclease free water. Total RNA sample quality was assessed via Agilent Bioanalyzer using the RNA 6000 Nano kit, resulting in RNA Integrity Scores (RIN) of 10, for all samples.

### Synthetic RNA Reference Standards

Spike-In RNA Variants (SIRVs) that include Iso Mix E0, ERCC, and Long SIRV modules were purchased from Lexogen (SIRV-Set 4, 141.03). SIRVs were resuspended in 10 µL Nuclease-free water to a concentration of 5.35 ng/µL. Resuspended SIRVs were then admixed into total RNA preparations prior to PolyA+ selection, CASPR treatment, or Reverse Transcription, to a concentration of 0.13 ng of SIRVs per µg of total RNA sample.

### Chemical Ablation of 3’ RNA ends

Sodium Periodate (NaIO_4_) was purchased from Millipore Sigma (311448-5G). A 2X Buffered Periodate Solution (BP) is prepared fresh each time by measuring NaIO_4_ powder and resuspending to a concentration of 4 mg/mL in aqueous solution of 200 mM Sodium Acetate pH 5.5 (Invitrogen, AM9740). Input RNA (up to 5 µg) is mixed with an equal volume of 2X BP and incubated at room temperature in the dark for 30 mins. Following treatment, RNA is cleaned with RNA RNA Clean & Concentrator (Zymo Research, R1013) according to the manufacturer’s instructions, and eluted in Nuclease Free Water.

### PolyA selection

Poly-adenylated transcripts were enriched from total RNA using the NEBNext Poly(A) mRNA Magnetic Isolation Module (E7490S), according to the manufacturer’s instructions.

### Reverse Transcription and Second Strand Synthesis

Reverse transcription is carried out with SuperScript IV Reverse Transcriptase (ThermoFisher, 18090010) or MarathonRT (Kerafast, EYU007). Reactions are carried out in a 20µL volume with the following components and final concentrations: 1X Reaction Buffer, dNTPs (0.5 mM), RNAseOUT (2U/µL), Oligo-d(T) primer (1 µM) or 4609bp gene specific primer (0.1 µM), 5 mM DTT (for SuperScript IV only), RNA input (<5 µg), and MarathonRT (0.5 µM or 20U) or SuperScript IV RT (200 U). Primers are initially annealed to template RNA in the presence of dNTPs, by heating to 65°C for 5 min, followed by snap cooling to 4°C for 2 mins. After snap cooling, the rest of the components are added, followed by reverse transcription for 1.5 hours at 42°C for MarathonRT and 50°C for SSIV. Reactions are stopped by heat inactivation at 85°C for 5 mins. Second strand synthesis is carried out using a modified Gubler and Hoffman procedure (26) adapted from Invitrogen’s A48570 kit, in a single pot format involving direct addition of second strand buffer, dNTPs, E.coli DNA Pol I, RNAse H, and E.coli DNA Ligase to the heat inactivated first strand reaction. Second-strand synthesis is carried out at 16°C for 2 hours, followed by DNA Clean with the Monarch kit for downstream processing. Verification of yield and quality of cDNA is determined via NanoDrop spectrometry, and by running on an 0.8% E-Gel NGS and imaged using Azure c600 (Azure biosystems)

### Nanopore Sequencing

All samples were barcoded with Native Barcoding kit (EXP-NBD104) prior to Nanopore library preparation using the Ligation Sequencing Kit (SQK-LSK109). All samples sequenced with MinION R9.4.1 flowcells, basecalled with Guppy basecaller 3.4.5, and demultiplexed with Guppy barcoder.

### Reference Sequences

A custom ribosomal RNA reference file was created by concatenating the fasta sequences for 28S (Gene ID: 100008589), 5.8S (Gene ID: 100008587), 5S (Gene ID: 100169751) and 18S (Gene ID: 100008588) ribosomal RNA sequences. lncRNA transcripts in fasta format were downloaded from Gencode release 31 (GRCh38.p12). For Human Reference alignment the UCSC analysis set of Dec. 2013 human genome (GCA_000001405.15) without the alt-scaffolds was used along with its associated gtf annotation file when appropriate. A custom reference sequence for R7 viral strain present in J-Lat cells was generated by extracting mapped reads from previous HIV alignments, size filtering, assembling with Unicycler (https://github.com/rrwick/Unicycler), polished with Medaka, and manually inspected with SnapGene against HXB2 originating background sequence to rule out structural variants.

### Determination of uniquely mapped reads

Reads were mapped to rRNA reference using *minimap2* with *map-ont* preset. Unmapped reads were extracted from the sam output using *samtools view* followed by conversion to fastq using *samtools bam2fq*(27). Fastq file containing unmapped rRNA reads were mapped to lncRNA reference with *minimap2* using *splice* preset, followed by extraction of unmapped reads and conversion to fastq as before. Unmapped lncRNA reads were remapped to human reference with *minimap2* using *splice* preset. Uniquely mapped reads were counted for each resulting sam file using *samtools view* with *-F260* flag to only count primary alignments and the *-c* option to output number of reads.

### Gene Body Coverage, Splice Junction Number, Read Distribution

For Gene Body Coverage calculation(28), reads were mapped directly to hg38 analysis set reference using *minimap2* with *splice* preset and *--secondary=no* flag, with mapped reads converted to bam format, sorted and indexed using *samtools*. Gene Body Coverage is calculated with the geneBody_coverage.py script that is part of the RSeQC package (v3.0.1) using sorted and indexed bam files and the UCSC RefSeq (refGene) annotations in bed format. Splice junction quantification and saturation was calculated using the *junction_saturation*.*py* script, also within RSeQC package, and with identical inputs as before. For Intragenic and Intergenic read distributions, reads were mapped and processed as before using the gencode v31 human reference (GRCh38.p12). The comprehensive genome annotation gtf file was collapsed using GTEx collapse annotation script. Read distributions were computed from mapped reads and collapsed annotations using RNA-SeQC (v2.3.4) with the following options *--unpaired --coverage –base-mismatch=180 --mapping-quality 0 --detection-threshold=0 -- legacy*.

### Statistical Analysis

Where indicated, t-tests were run between CASPR and PolyA-selected samples within RT enzyme group (either MRT or SSIV). Analyses performed within GraphPad Prism 8, assuming all rows are sampled from populations with same scatter (SD). Statistical significance determined using the Holm-Sidak method, with alpha = 0.05. Statistical significance denoted as following: p<0.05 (*), p<0.01 (**), p<0.001 (***), p<0.0001 (****)

### HIV isoform analysis

Reads were mapped to R7 reference sequence with *minimap2* using *splice* preset, followed by filtering using *-F260* flag in *samtools view* and sorting. Resulting sorted bam file is used as input for Pinfish pipeline (https://github.com/nanoporetech/pinfish). Briefly bam files were used as input for *spliced_bam2gff* command using the *-M* option. The resulting gff file is clustered into isoform bins using *cluster_gff* command using the following options *-c 3 -p 0*. Isoforms clusters are then polished using polish_clusters command with *-c 3* option. Polished clusters in fasta format are remapped to reference using minimap2 and processed using same settings as before. Polished clusters are visualized at this stage using IGV 2.7.2, and coverage maps for clustered isoforms are obtained with the *samtools depth* command with the *-a -d 0* options. The *spliced_bam2gff* command is then run with identical options as before and resulting polished clusters that are then collapsed with the *collapse_partials* command with the *-M -U* options. Resulting GFF files are then manually parsed to retain those isoforms that contain at least one full CDS.

### Host cell transcript isoform analysis

Analysis of host cell isoforms was performed using the FLAIR pipeline (29) v1.4. Reads are mapped to UCSC hg38 reference using *flair align* module using option *-p*, followed by splice junction correction with the *flair correct* module. Isoforms are collapsed using the *flair collapse* module with *--stringent --trust_ends* options to ensure 80% coverage per isoform cluster. Transcript lengths can be calculated with flair collapse outputs, by indexing the transcripts.fa file for each sample with *samtools faidx* and extracting the second column containing length of each sequence. The isoforms are then quantified with the *flair quantify* module using *--tpm --trust_ends* options. Outputs of this module were used to compute gene expression TPM correlation between samples and replicates. The *flair diffexp* module is finally used to generate differential gene/isoform expression analysis with default settings. Finally, the *flair diffsplice* module is used to determinate high confidence alternative splicing events from the isoforms processed with previous modules. Differential gene, isoform or splicing outputs are filtered for max p-value of 0.1, those hits that remain are subject to additional FDR analysis with those with p-adj<0.1 being highly significant. Transcript discovery sensitivity and specificity was calculated using *gffcompare* v0.11.5(30) using gtf files outputs from *flair collapse* module and the UCSC hg38 genome annotation in gtf format with the following command options *-T -M -r*.

### Cross validation with published Illumina dataset that used J-Lat 10.6 clone and TNF-alpha induction

Raw Illumina PE150 data from Ma et al. 2021 manuscript (31) was downloaded from the SRA/ENA repository, with two replicates obtained for each TNF-plus and TNF-minus treatments (Accessions: SRR13649912, SRR13649913, SRR13649916, SRR13649917). For analysis of host transcripts, reads were mapped to the hg38 reference using HISAT2 with the following settings *--rna-strandness RF*, and converted to a sorted bam file with samtools. Human reference mapping counts per replicate were used to normalize raw transcript count matrix obtained for this dataset from the Gene Expression Omnibus (Accesion: GSE166337), yielding Counts Per Million (CPM) metric. Genes shown to be significant in our Nanopore DGE data (p-adj<0.1) were queried against normalized counts from PE150 dataset to determine concordance in fold-change direction before and after TNF-alpha induction. For splicing site usage analysis, sorted bam files were processed with Portcullis (32) with the following settings *--exon-gff --orientation RF --strandedness firststrand --separate* to obtain spliced/unspliced bam and gff files. Splice site ROIs are extracted from gff file, and normalized to obtain CPM values for a given splice junction. For analysis of HIV transcripts from this short read data, reads were mapped to the R7 HIV reference sequence with HISAT2 with the following settings *--rna-strandness RF --max-intronlen 5000*, followed by conversion to sorted bam file using samtools. Sorted bam files were processed with Portcullis with same settings as before to obtain spliced and unspliced bam and gff files. Coverage of spliced HIV reads was computed using samtools depth with the following settings *-a -d 0*, with resulting depth per position normalized to average number of reads for a given dataset (to allow comparison with other datasets).

### DGE subset validation via Relative Quantification qPCR

Total RNA was isolated from J-Lat 10.6 cells treated with TNF-alpha (n=3) or vehicle control (n=4), with RNA integrity verified with Agilent Bioanalyzer RNA Nano chip. Total RNA was treated with CASPR and Reverse transcription was carried out with MarathonRT and Oligo-d(T) priming, followed by second-strand synthesis with identical conditions to those use in sequencing samples. Double stranded cDNA purity and concentration was assessed with Nanodrop 1000, followed by sample dilution to 0.1-0.5 ng/µL. PrimeTime qPCR primers were ordered from Integrated DNA technologies for the following gene targets:

- ACTB, RefSeq NM_001101, Product ID Hs.PT.39a.22214847: 5’-ACAGAGCCTCGCCTTTG-3’, 5’-CCTTGCACATGCCGGAG-3’ (Exon 1-2)
- LIMD2, RefSeq NM_030576, Product ID Hs.PT.58.25965347.g: 5’-CACTGTCACACCAAGCTCAG-3’, 5’-GTCGTAGTTGCCTTTGCTCTT-3’
- MYO7B, RefSeq NM_001080527, Product ID Hs.PT.58.1328863: 5’-CCGAATCCAGAAGGTCCTGA-3’, 5’-CAACCACCTCCAGCATCTC-3’ (Exon 38-40)
- PSAT1, RefSeq NM_058179, Product ID Hs.PT.58.20540177: 5’-GTCCTCAAACTTCCTGTCCAA-3’, 5’-TCATCACGGACAATCACCAC-3’ (Exon 5-6)
- TNFAIP3, RefSeq NM_006290, Product ID Hs.PT.58.1824217: 5’-TGATAGAAATCCCCGTCCAAG-3’, 5’-TCCTGCCATTTCTTGTACTCAT-3’ (Exon 6-7)

qPCR was carried out using 5 µL of cDNA input, 500 nM primers, and Luna Universal qPCR Master Mix at 1X concentration (NEB, M3003L) in total reaction volume of 20 µL. Plates were run in a Roche Lightcycler II instrument with the following cycling parameters with ramp rates of 2.2 °C/s: 95°C for 1 min (Initial Denaturation), 45 cycles of 95°C for 15 sec and 60°C for 30 sec (Amplification), 95°C for 5 sec and 60°C ramp to 95°C (Melting Curve), 40°C for 10sec (Cooling). qPCR was validated by running a serial dilution of Jurkat total RNA across 6-logs to determine PCR efficiency, and Standard curve Slope, Y-intercept, and R2 error for each primer set: ACTB (1.799, -3.921, 16.91, 0.0273), LIMD2 (1.927, -3.512, 21.00, 0.00429), MYO7B (1.916, -3.541. 18.02, 0.0133), PSAT1 (1.937, -3.482, 19.02, 0.0158), TNFAIP3 (1.980, -3.371, 22.91, 0.0579). Non-Template Controls were used for all reactions, with Ct values >40 cycles for all samples. Gene expression was calculated via Relative Quantification of LIMD2, MYO7B, PSAT1 and TNFAIP3 Targets over ACTB Reference, with two technical replicates used per sample. Gene expression for +/-TNF-alpha samples was determined using the Adv. Relative Quantification Module in Lightcycler 480 Software (version 1.5.0 SP3) in High Confidence Mode.

## Results

### Improvement of the specificity and yield of high performing reverse transcriptases for producing full-length transcripts for direct cDNA sequencing

Obtaining a readout of alternative splicing of host and viral transcripts involves end-to-end sequencing reads which provides for full-exon connectivity. To achieve this, processive reverse transcriptases are required, along with an enrichment scheme to select for protein coding sequences (CDS) from total RNA isolates. For direct cDNA sequencing, an additional requirement is to maximize the yield of cDNA so as to dispense with the need for PCR amplification of transcripts. Taking into account these requirements, we first evaluated the high performing reverse transcriptases MarathonRT (MRT), a eubacterial group II intron that has been shown to efficiently copy structured long RNAs (33,34), and SuperScript IV (SSIV), which has been considered a “commercial gold standard” (35,36), for their yield of protein coding transcripts from Nalm6 total RNA, a human leukemic B cell line.

Gel electrophoresis of double-stranded cDNA obtained via SSIV and MRT showed prominent bands of similar size to ribosomal RNA when Nalm6 total RNA is directly reverse transcribed with Oligo-d(T) priming without any CDS enrichment strategy (i.e. Control) **(Figure 1A)**. The presence of putative rRNA bands when using total RNA was unsurprising given these structural RNAs are a major RNA cellular component (enriched up to 90% in total RNA) and source of interference in RNA-Seq workflows (37,38). However, ribosomal RNAs are not polyadenylated, which raises the question on the source of this spurious priming. We hypothesized these primer-independent products were the result of the RNAs themselves priming the RT initiation complexes, and that blocking 3’-OH ends of RNA inputs prior to reverse transcription could be beneficial in increasing the specificity of RT priming. For this purpose, we developed an approach dubbed **C**hemical **A**blation of **S**puriously **P**riming **R**NAs (CASPR) based on the oxidation of vicinal 2’- and 3’-OH diols of RNA, which results in the ablation of 3’-OH ends only in RNA, preventing their priming during RT and favoring RT initiation from the intended exogenous Oligo-d(T) DNA primer. Pre-treatment of input RNA with CASPR visibly improved RT specificity in both SSIV and MRT, resulting in a smear reminiscent of PolyA+ selection (PolyA+) **(Figure 1A)**, albeit with greater mass yield compared to this established methodology **(Figure 1B)**. The increases in specificity of Oligo-d(T) priming elicited by CASPR treatment were particularly evident in MRT samples, where CASPR treated lanes do not show any discernable rRNA bands, compared to residual rRNA bands present with SSIV. CASPR treatment also resulted in 5- and 10-fold improvements in cDNA yield compared to PolyA+ for SSIV and MRT respectively (p<0.01 and p<0.001), with the CASPR MRT combination resulting in 50% greater cDNA yield compared to CASPR SSIV (p<0.05). This increase in RT specificity was consistent when using total RNA from other human cell lines, and when using gene-specific priming modalities with *in vitro* transcribed HIV RNA **(Supplementary Figure 1)**, suggesting spurious priming from RNA inputs is prevalent. Importantly, CASPR treatment did not compromise RNA quality as measured on total RNA via Agilent Bioanalyzer, with RNA integrity (RIN) scores of 10 obtained from both CASPR and vehicle controls **(Supplementary Figure 2)**.

**Figure 1.**
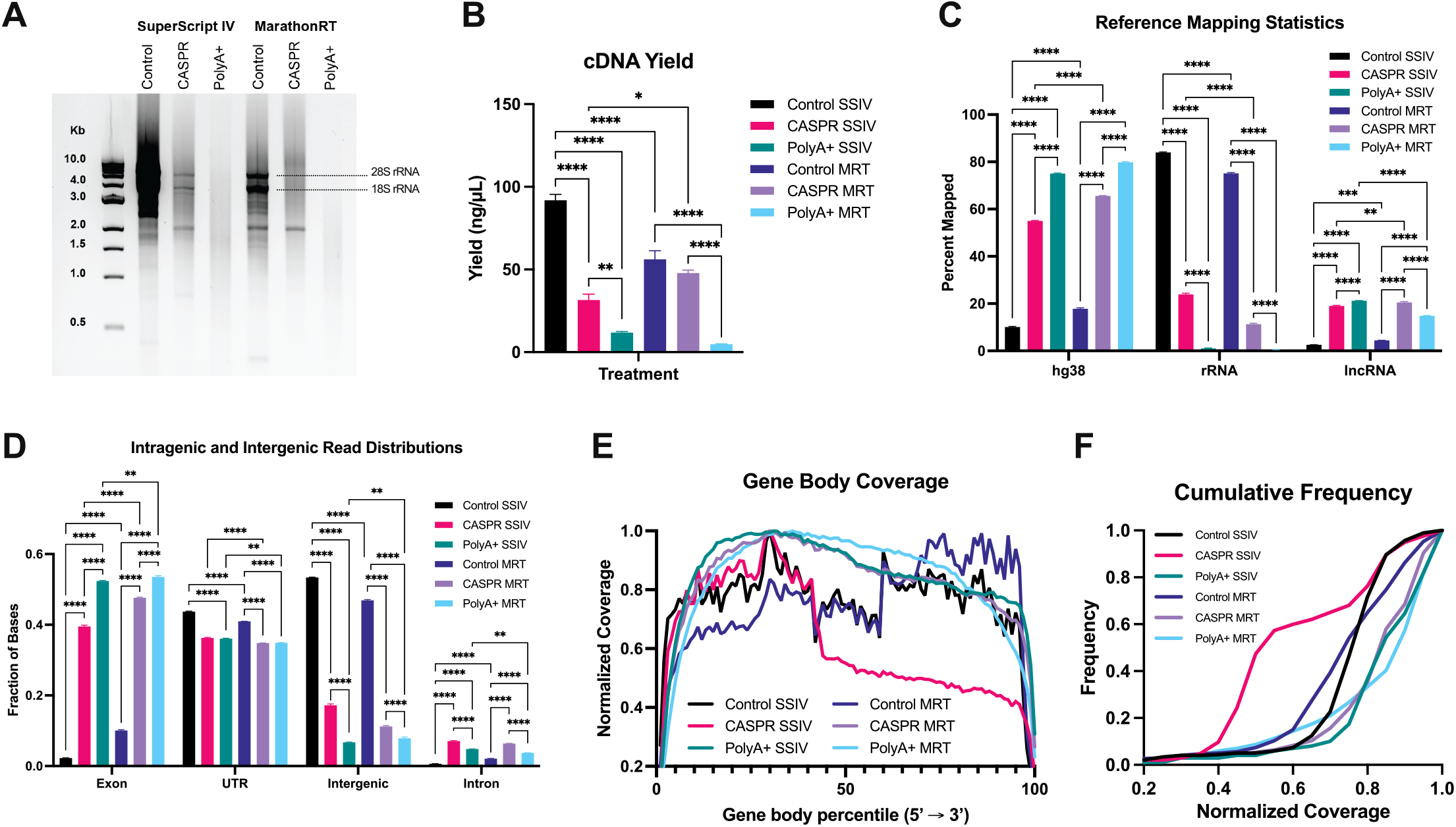
CASPR improves the specificity of Oligo-d(T) primed RT when using total RNA inputs by reducing rRNA and increasing coverage evenness of protein-coding transcripts. **(A)** 1% Agarose gel electrophoresis of double stranded cDNA products that were Reverse Transcribed with Oligo-d(T) priming with SSIV or MRT with no CDS enrichment (Control), CASPR or PolyA+ selection. **(B)** cDNA yield of different RT and CDS enrichment combinations as measured spectrophotometrically. **(C)** Fraction of reads uniquely mapped to the listed references using Nanopore sequencing. **(D)** Intragenic and Intergenic reads distributions. **(E)** Gene body coverage of protein coding transcripts, and **(F)** Cumulative Frequency distribution of Gene Body Coverage. All values are means ± SEM. Statistical significance calculated with two-way ANOVA with Tukey multiple comparisons test, p<0.05(*), p<0.01(**), p<0.001(***), p<0.0001(****)

To validate that CASPR treatment was reducing rRNA, cDNA samples were sequenced with Oxford Nanopore Technologies (ONT) MinION to determine the effect of CASPR treatment at the read mapping level **(Figure 1C)**. As expected, the most prominent effect of CASPR treatment is the reduction of reads mapping to rRNA reference from 84% to 24% in SSIV and from 75% to 12% for MRT respectively (p<0.0001 for both). This reduction in rRNA-mapped reads in CASPR-treated samples was associated with a proportional increase in percent of reads mapping to the human genome (hg38) reference, from 10% to 55% in SSIV and from 18% to 66% in MRT (p<0.0001 for both), which compares favorably with hg38 enrichment levels in PolyA+ samples (75-80%). Compared to PolyA+, reads mapped to lncRNA mapping fractions were mostly nominal after CASPR treatment in both SSIV and MRT samples. Despite substantial CASPR-elicited increases in Oligo-d(T) priming specificity, the reductions in rRNA were not fully penetrant compared to PolyA+, which routinely reduced rRNA reads to ∼1% irrespective of RT used. However, the read mapping fractions also show that each RT is not equally susceptible to rRNA interference, with MRT showing 2-fold lower rRNA fractions and 20% higher hg38 fractions after CASPR treatment compared to SSIV, suggesting MRT is more amenable to the priming specificity improvements elicited by ablation of 3’ RNA ends when using total RNA inputs. Given improvements observed in mapped read distributions elicited by CASPR, we next evaluated its effect on the distribution of intergenic and intragenic reads **(Figure 1D)**. As expected, the most notable effect of CASPR and PolyA+ was a dramatic reduction in intergenic reads, with an associated increase in proportion of reads mapping to exonic and intronic regions (p<0.0001 for all comparisons). Interestingly, both CASPR and PolyA+ slightly reduced read mappings to UTR regions in both RTs despite the associated increases in exonic reads that were observed for either treatment. All of this points to largely equivalent effects of CASPR and PolyA+ in increasing proportion of reads mapping to the intragenic features that delineate exon connectivity.

In addition to mapping statistics, the coverage along the length of protein coding transcripts is critical to reveal full exon connectivity. For this purpose, hg38 mapped reads were cross-referenced with the RefSeq genome annotation file to delineate the coverage along the 5’ to 3’ axis of each expressed transcript, an approach known as gene body coverage (28). The gene body coverage when using total RNA without CDS enrichment shows inconsistent coverage, with the Control SSIV samples having clear 5’ and 3’ end biases (and associated low coverage in middle region of gene body), and Control MRT showing consistent 3’ end bias **(Figure 1E)**. Conversely, PolyA+ samples show even coverage with normalized coverage values consistently not dipping below 0.8 across 70% of gene body for both SSIV and MRT **(Figure 1F)** and no appreciable 5’ or 3’ end biases. These data suggest that the reduction in ribosomal RNA reads does not only increase the number of reads mapping to protein-coding transcripts, but also increases their evenness of coverage across the length of the transcript. Interestingly, MRT samples that are CASPR treated show a gene body coverage distribution similar to that observed in PolyA+ samples, also consistently above 0.8 normalized coverage across majority of transcript body **(Figure 1F)**. However, this same effect is not observed with CASPR-treated SSIV samples, with 0.5 median coverage compared to the >0.7 coverage values observed for all other treatment and RT combinations **(Figure 1F)**. Overall, this data underscores the importance of CDS enrichment strategies and processive RTs in obtaining full-exon connectivity, while highlighting potential benefits of CASPR as an alternative to PolyA+ selection to substantially increase RT yield and priming specificity when using total RNA inputs.

### Analytical Performance Validation of CDS-enrichment strategies and RT conditions using synthetic RNA reference standards

Initial optimization of RT conditions using processive enzymes and a novel CDS enrichment strategy suggests that the combination of MarathonRT with CASPR is well suited for direct cDNA sequencing using ONT. However, despite compelling data showing CASPR as a higher-yield analogue of PolyA+ selection, and the coverage improvements elicited with MarathonRT, neither of these interventions has been formally validated with reference standards. Synthetic RNA reference standards, which include ERCCs, SIRVs, and Sequins, have recently emerged for validating full RNA-Seq workflows (39), and contain synthetic poly-adenylated mono- and/or multi-exonic transcripts of varied characteristics and in known concentration ranges. Given the synthetic nature of these transcripts, resulting reads obtained via sequencing can be cross-referenced with ground-truth annotations to evaluate quantitative features of our workflow, the sensitivity and breadth of transcript capture, length biases due to RT processivity constraints, and other performance variables. We used a Spike-In RNA Variants (SIRV-Set 4) mix that was spiked into Nalm6 total RNA isolations prior to any enrichment interventions or RT with the goal of validating analytical performance of MarathonRT and CASPR against established gold standards in the field. Three different CDS enrichment strategies (Control, CASPR, PolyA+), and two different RTs (SSIV, MRT) were tested in triplicate using the same SIRV-spiked total RNA sample and run in parallel (**Figure 2A**). All the resulting CDS enrichment and RT combinations (6 combinations, with 3 replicates each) are thus technical replicates of each other, and sequenced and analyzed (**Figure 2B)** in parallel to allow for robust comparisons between each condition.

**Figure 2.**
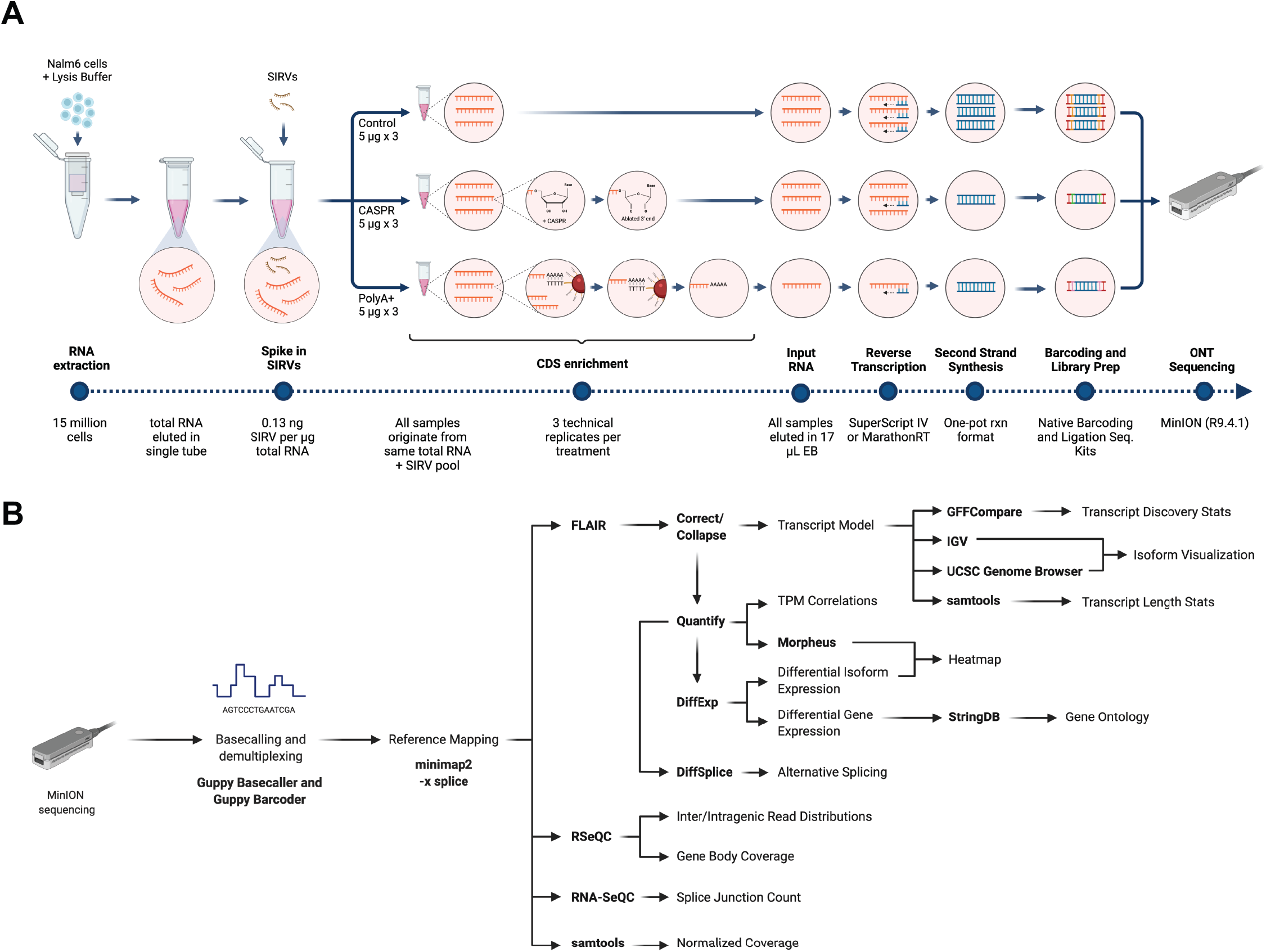
Assay and Bioinformatic workflow for analytical performance validation. **(A)** Total RNA was isolated from Nalm6 cells and pooled into a single tube. Total RNA was spiked with SIRV-Set 4 at a concentration of 0.13 ng SIRVs per microgram of total RNA. Three CDS enrichment conditions (Control, CASPR, and PolyA+ selection) were tested in parallel, all drawing identical RNA inputs from same total RNA sample in triplicate per condition. Following CDS enrichment, all samples are eluted in 17 µL of EB, followed by Reverse Transcription using 10 µL of input, and one-pot second strand synthesis using a modified Gubler and Hoffman method. Following second-strand synthesis cleanup, identical volumes of double-stranded cDNA samples are then barcoded and prepared for sequencing using Oxford Nanopore Technologies (ONT) Native Barcoding (EXP-NBD104, EXP-NBD-114), and Ligation Sequencing (SQK-LSK-109) Kits. Following library prep, all samples are eluted in identical volumes of EB, then equal volumes of each sample are pooled and sequenced via ONT MinION, using R9.4.1. chemistry. (B) Bioinformatic workflow used for data generation and analysis throughout manuscript. Typical outputs for each analysis are shown, with text in **bold** denoting specific tools used for a particular output.

Consistent with previous findings, direct cDNA sequencing of SIRV-spiked Nalm6 showed that CDS enrichment strategies are critical for enrichment of poly-adenylated synthetic transcripts **(Figure 3A)**. Specifically, CASPR treatment of total RNA prior to RT increased SIRV mapping by 5-fold in SSIV and 2.5-fold in MRT (p<0.001 and p<0.01 respectively). Moreover, the enrichment of SIRV reads with CASPR was comparable with that of PolyA+, with differences between enrichment strategies for each RT not statistically significant for SSIV and modest for MRT (p<0.05). Given the lack of meaningful SIRV mapping fractions without CDS enrichment, CASPR and PolyA+ samples were sequenced deeper to allow for more sensitive analysis. One such analysis involves the quantification of ERCC controls within the SIRV mix, which are present in known concentrations spanning 6-logs. Cross-referencing of measured expression of ERCC transcripts with known input amounts, showed that cDNA measurements are quantitative, with R^2^ values averaging 0.9 for all CDS enrichment strategies and RT combinations **(Figure 3B)**. This robustness in cDNA quantitation translates to actual measurements of human transcript abundance with all TPM correlations strongly trending in a linear manner irrespective of RT or CDS enrichment strategy tested **(Figure 3C)**. ERCC data and hg38 gene expression correlations are strongly suggestive of CASPR treatment being functionally equivalent to PolyA+ selection with regards to ability to accurately quantify cDNA levels despite residual rRNA and marginally lower hg38 mapping fractions **(Figure 1C)**. However, this does not provide clarity on the extent of coverage of these transcripts, a critical variable for full-length sequencing.

**Figure 3.**
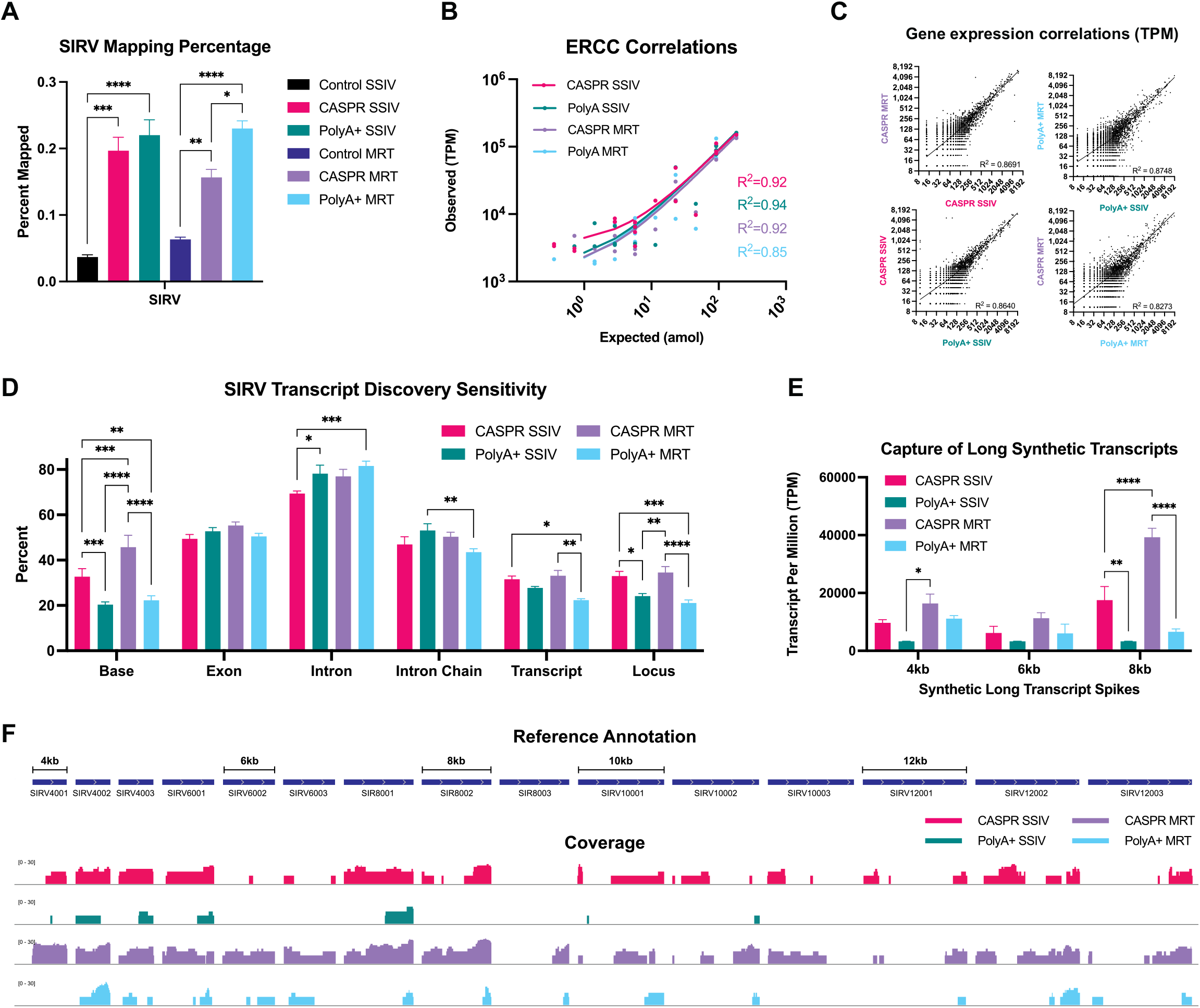
Validation with synthetic reference standards show CASPR is functionally equivalent to PolyA+ selection, but results in higher cDNA yield, coverage evenness and capture of long transcripts. **(A)** Percent of reads uniquely mapped to SIRV reference sequences. **(B)** Correlations of gene expression TPM values with absolute input amounts of each synthetic transcript (in attomoles) for ERCC subsets. **(C)** Hg38 gene expression correlations between different RTs and CDS enrichment strategies. **(D)** Transcript Discovery Sensitivity calculation using FLAIR-derived transcriptome and hg38 gtf annotation file. **(E)** Efficiency of capture of Long SIRVs of 4kb, 6kb, and 8kb size classes. **(F)** Raw coverage visualized via IGV of all Long SIRVs for each RT and CDS enrichment strategy combination. All samples ran in triplicate (n=3). All values are Means ± SEM. Statistical significance calculated with two-way ANOVA with Tukey multiple comparisons test, p<0.05(*), p<0.01(**), p<0.001(***), p<0.0001(****)

Isoform-level analysis can add an additional layer on the breadth of transcript coverage elicited by different RTs and CDS enrichment strategies. Isoform collapse and quantification of SIRV transcripts using FLAIR (29), followed by cross-referencing to known SIRVome reference annotation files shows that transcript capture sensitivities are largely equivalent between CASPR and PolyA+; however, CASPR provides distinct improvements in the transcript discovery sensitivity at the Base Level **(Figure 3D)**. Base level transcript discovery corresponds to the number of exon bases within a query transcript that are reported at the same coordinate as the reference annotation. Therefore, a higher base level number would be reflective of a longer section of sequenced transcripts overlapping the reference annotation, and indicative of a greater breadth of coverage (30). In this regard, CASPR treatment shows 2-fold higher transcript discovery sensitivity at the Base level compared to PolyA with both SSIV and MRT (p<0.001 and p<0.0001 respectively) and 40-60% higher at the Locus level (p<0.05 for SSIV, p<0.0001 for MRT). This suggest that even though CASPR and PolyA result in equivalent number of read counts per transcript, CASPR provides significantly higher coverage per captured transcript, resulting in increased practical throughput and higher likelihood of capturing full-exon connectivity. Finally, Long SIRVs ranging from 4-12 kb were quantified after sequencing for all RTs and CDS enrichment conditions to evaluate the propensity of each treatment combination to result in size biases related to the inherent processivity constrains of RTs for RNA inputs greater than 5 kb in length, which was previously reported by us (33). Compared to PolyA+, CASPR trended toward increased sensitivity for capture of long synthetic transcripts greater than 5 kb in size for all size classes **(Figure 3E)**. Of particular note is the statistically significant increase in sensitivity of capture of 8kb transcripts elicited by CASPR treatment, resulting in 6-fold increases in capture for both SSIV and MRT (p<0.01 and p<0.0001 respectively), and with MRT resulting in 2-fold higher sensitivity for this transcript size class as compared to SSIV (p<0.0001). This increase in sensitivity of capture in CASPR treated samples also translated to increased breadth of coverage for all transcript classes, with MRT in combination with CASPR showing more even coverage across all Long SIRV transcript size classes, compared to limited coverage obtained with PolyA+ for both RTs **(Figure 3F)**. Overall, this data validates CASPR is functionally equivalent to PolyA+ selection while providing distinct advantages such as greater transcript coverage sensitivity and greater capacity to capture long transcripts. In addition, this data confirms that MarathonRT, in combination with CASPR, has superior sensitivity and breadth of coverage than SSIV for capturing long polyadenylated transcripts from complex mixtures of host cell mRNAs.

### Evaluation of RT and CDS enrichment strategies in the J-Lat 10.6 T cell line undergoing active HIV transcription

To determine whether our direct cDNA sequencing workflow can effectively capture HIV RNAs within a swarm of host cell transcripts, we evaluated both RTs and CDS enrichment conditions using the J-Lat 10.6 lymphocytic CD4 T cell line(23). This established and well-characterized Jurkat cell line has a single integrated provirus that contains all canonical splice sites and can be robustly induced to produce viral RNAs with TNF-alpha or other suitable HIV reactivation agents (24). Moreover, activation results in production of physiological levels of viral RNA, while also being representative of host transcriptional regulation dynamics of active infection(23). Thus, the J-Lat 10.6 cell line provides a stringent test case for evaluating efficiency of viral isoform capture within dynamically changing host cell transcripts without relying on PCR amplification to enrich for rare transcript variants, while allowing us to examine the effects of HIV reactivation on host cell transcript regulation. J-Lat 10.6 cells were induced with 10 ng/mL TNF-alpha for 24 hours, followed by assessment of p24 induction and EGFP expression, with all induction values normative to previous publications **(Supplementary Figure 3)**. Both SSIV and MRT were tested for their performance with CASPR or PolyA selection, with all replicates and samples run in parallel. As consistent with previous data, host cell gene expression TPM values show concordance between CASPR treatment and PolyA+ selection when using either SSIV or MRT **(Figure 4A)** and was reproducible across replicates **(Supplementary Figure 4)**. Normalized gene body coverage values are consistent with those found in Nalm6 datasets, with CASPR MRT samples approaching the evenness observed in PolyA+ selected samples, and with SSIV showing measurable 5’ end bias as consistent with previous data **(Figure 4B)**. Compared to PolyA+, CASPR increases the fraction of long transcripts >4000 bp by 2.5-fold and 6-fold in SSIV and MRT respectively (p<0.05 for both), in a manner that is consistent with previously observed enrichments of Long SIRVs **(Figure 4C)**. The ability to capture longer transcripts positions CASPR well for the capture of HIV transcripts which are intrinsically difficult to reverse transcribe given high RNA structure (40) and their relatively long length (2-4 kb for spliced viral transcripts) compared to host cell coding transcripts (∼1kb average size).

**Figure 4.**
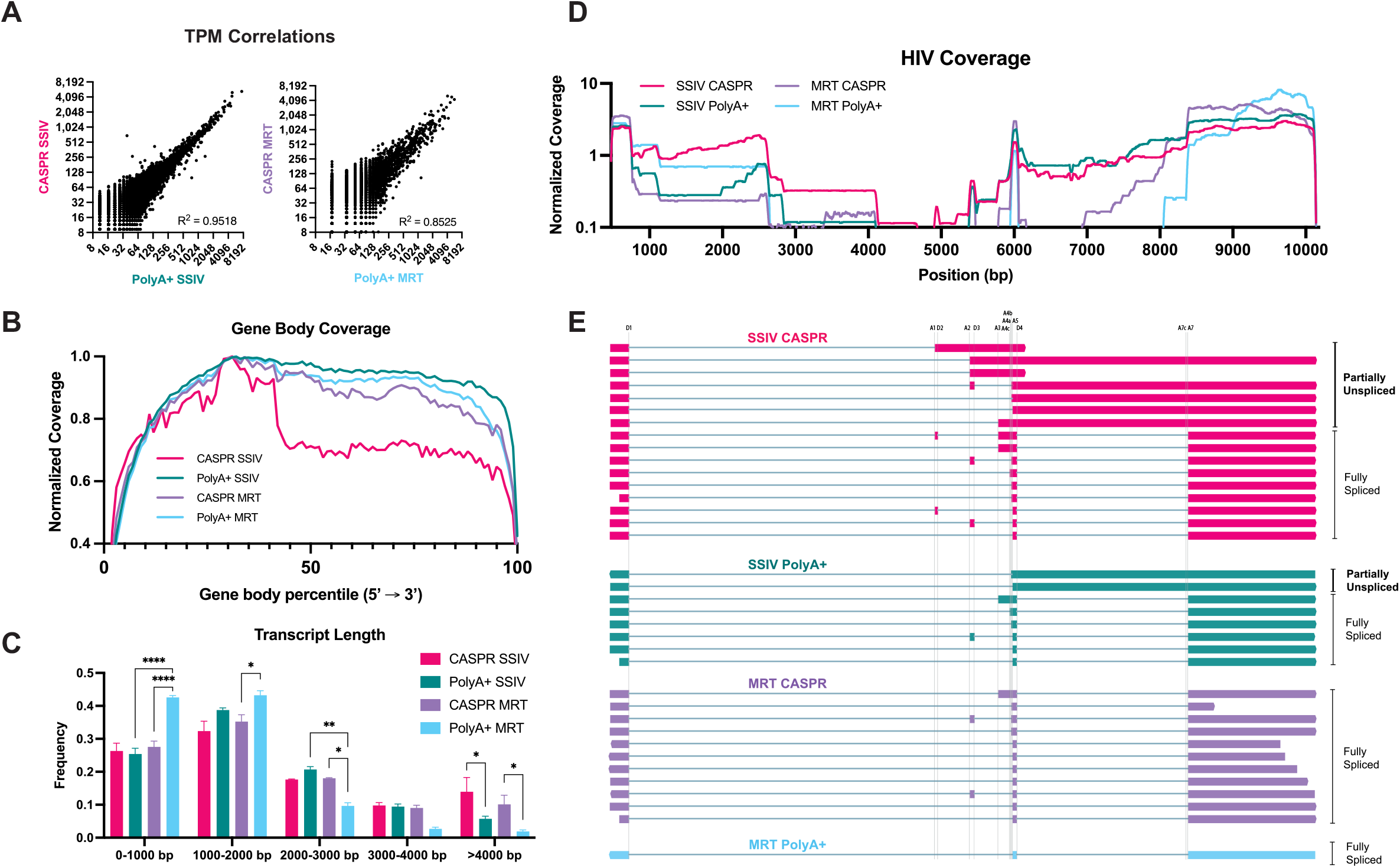
Evaluation of RT conditions and CDS enrichment strategies in capture of host and viral transcripts in cell line actively expressing HIV. **(A)** Host cell gene expression correlations for each CDS enrichment strategy when using SSIV and MRT. **(B)** Gene-Body coverage of protein coding hg38 transcripts. **(C)** Frequency of host cell transcript lengths derived from FLAIR isoform analysis pipeline, binned at 1000bp intervals. **(D)** Coverage Map of raw reads. All samples ran in duplicate (n=2). **(E)** Visualization of isoform structure of multiexonic HIV transcripts processed with Pinfish pipeline. All values are means ± SEM. Statistical significance calculated with two-way ANOVA with Tukey multiple comparisons test, p<0.05(*), p<0.01(**), p<0.001(***), p<0.0001(****)

With regards to the capture efficiency of HIV transcripts, our pipeline was able to capture thousands of HIV reads despite constituting less than 1% of total dataset **(Supplementary Figure 5)**. To compare the performance of RTs and CDS enrichment strategies in coverage evenness, reads were mapped to the HIV reference and normalized across length of the genome, with a normalized coverage of 1 indicating even sampling **(Figure 3D)**. CASPR and SSIV shows more consistent coverage across length of genome, with relative coverage being close 1 for most of the genome tract length relevant to multiexonic transcripts (5,000-10,000bp). SSIV PolyA trails closely behind, but shows reduced coverage in regions associated with Vif and Vpr transcripts (5000-6000bp), and overall lower coverage for regions coding for Gag and Gag-Pol. Compared to SSIV, MRT shows 3’ end bias and with coverage dropping between 7500-8300 bp. Of particular note, all samples show sharp increases in coverage at ∼2700 and ∼4200 bp which are inconsistent with any splice junctions. However, the presence of long poly-adenylated stretches in these two regions are suggestive of mispriming of Oligo-d(T) being responsible for these artefactual increases in coverage.

To evaluate HIV isoform diversity in all treatments, HIV-mapped reads were grouped by exon boundaries into isoform clusters and collapsed into high confidence multiexonic transcript models. This analysis pipeline worked robustly and identified splice sites that were consistent with those previously observed with long-read sequencing approaches **(Supplementary Table 1)**. Multiexonic transcripts identified by Pinfish were then parsed to determine likely expressed gene based on which undisrupted open reading frame (ORF) is closest to the 5’ end **(Figure 3E)**. As consistent with normalized coverage data, MRT with PolyA+ selection did not capture overall HIV isoform diversity, with fully-spliced species being favored. MRT with CASPR treatment performs nominally better than PolyA selection in increasing the isoform diversity of fully spliced transcripts; however, this treatment combination does not capture any partially unspliced transcripts coding for Env, Vpr and Vif. SSIV in combination with CASPR shows overall highest HIV isoform diversity, resulting in an assortment of fully-spliced transcripts and 2-3 fold higher capture of partially spliced species compared to PolyA+. The detectable differences in viral isoform diversity captured with MRT and SSIV highlight the need to evaluate each RT enzyme independently of their performance in the capture of host cell transcripts and adopt strategies that take advantage of each RT’s unique characteristics and strengths. For this purpose, an optimized approach to increase the likelihood of capturing *<u>both</u>* host and viral samples would rely on the interrogation of CASPR-treated total RNA using both SSIV and MRT, followed by the simultaneous sequencing of resulting cDNA.

### Differential expression analysis using optimized RT and CDS enrichment conditions identifies alternatively-spliced host factors of HIV assembly and defines their associated HIV splicing signature

Having evaluated the role of CASPR in increasing transcript capture efficiency and coverage metrics, and the identified strengths of SSIV and MRT for capture of respective viral and host transcripts, we set out to do a larger scale survey of viral reactivation dynamics within host cells in the J-Lat 10.6 cell line. The goal was the simultaneous identification of differentially regulated transcripts within host cells and their HIV isoform correlates. Taking into account our previous findings regarding the unique suitability for SSIV and MRT in the efficient capture of respective viral and host transcripts, total RNA was treated with CASPR and then split evenly to be reverse transcribed with SSIV and MRT, with resulting cDNA being used for sequencing. Since TNF-alpha induction is likely to cause global perturbations in host cell gene expression, the effect of TNF-alpha in J-Lat 10.6 case group was compared with the differentially regulated transcripts elicited by TNF-alpha treatment in a control group of parental Jurkat cells lacking an integrated provirus. Those transcripts found to be differentially regulated by TNF-alpha in Jurkat control group, was ‘subtracted’ out from those differentially regulated in J-Lat 10.6 case group, which will provide greater clarity on the host-cell transcripts that will be uniquely up/down regulated by active HIV transcription, and not by the HIV reactivation agent itself.

An initial pilot run showed suitability of the approach in using both MRT and SSIV to maximize respective host cell and viral transcript capture efficiencies and coverage breadth during sequencing. Specifically, MRT showed 4-fold lower capture of artefactual rRNA-related hits in pilot differential isoform expression (DIE) analysis as compared with SSIV, with the latter showing ∼40% of DIE hits can be traced to rRNA loci **(Supplementary Figure 6)**. Given these initial results confirming suitability of our split MRT/SSIV approach, we proceeded to sequence additional biological replicates (up to a total of 5) in the presence or absence of TNF-alpha for both J-Lat (Case) and Jurkat (Control) groups **(Supplementary Table 2)**. Differential gene expression (DGE) analysis upon TNF-alpha induction in both case and control groups with (p-values<0.1), revealed 244 and 139 genes passed this filtering criteria in J-Lat case and Jurkat control groups respectively **(Supplementary Figure 7)**. Of those genes passing p-value filtering criteria, 20 genes were found to be modulated by TNF-alpha induction in both J-Lat and Jurkat datasets, suggesting relatively low overlap between responses to TNF-alpha induction in Case and Control groups. To further determine the extent of TNF-alpha response overlap between case and control groups, DGE data was used to compute functional enrichment analysis with StringDB(41) version 11.0 with Gene Ontology (GO) framework at the Cellular Component and Biological Process levels. Highly significant (FDR<0.01) GO Cellular components enriched in J-Lat case group include the ‘NF-kappaB complex’, the ‘spliceosomal complex’, and ‘secretory granule membrane’, which do not overlap with the single ‘cytosolic ribosome’ term found in Jurkat control group **(Table 1)**. Likewise, GO Biological Process terms do not overlap between case and control groups, except for ‘NF-kappaB signaling which is present in both case and controls groups but 2-fold more enriched in the former group **(Supplementary Figure 8)**. The activation of the NFKB complex observed in functional enrichment analysis is consistent with the highly significant (p-adj < 0.05) genes found to be differentially regulated in J-Lat cells treated with TNF-alpha including: TNFAIP3, NFKBIA, BIRC2, and NFKB2 **(Table 2)**. Cross-comparison of our highly significant DGE hits (p-adj<0.1) with a publicly-available NGS dataset from a report using the J-Lat 10.6 clone and similar TNF-alpha induction conditions shows high concordance, with 78% of genes showing a consistent and highly significant fold-change direction in comparison with our data after TNF-alpha induction (**Supplementary Figure 9**) (31). Moreover, we performed qPCR validation of a subset of significant DGE hits and show high concordance with our sequencing findings both in the direction of fold changes upon TNF-alpha induction and their statistical significance **(Supplementary Figure 10)**. A significant fraction of the DGE hits in the J-Lat case group (including those related to NFKB complex) were also found to be highly significant in Jurkat group, underscoring the utility of our ‘subtractive’ approach to tease apart partially overlapping responses. The NFKB complex related genes that were found to be differentially expressed exclusively in J-Lat cells include NFKBIA and BIRC2, which were previously found via RNA-Seq to be upregulated upon latency reversal in SIV-infected ART-suppressed non-human primates (42). BIRC2 was also found to be a negative regulator of HIV-transcription that could be antagonized with Smac mimetics for reversal of latency (43). The robust upregulation of BIRC2 we observed in our data set despite active HIV-transcription, can be reconciled with the paradoxical role of this gene as both a positive modulator of the canonical NFKB (cNFKB) pathway and a negative modulator of the non-canonical NFKB (ncNFKB) pathway (44), with our use of TNF-alpha engaging the cNFKB pathway.

**Table 1.** Significant functional enrichments elicited by TNF-alpha.

| #term ID | term description | genes mapped | enrichment score | FDR | method | Sample Group |
| --- | --- | --- | --- | --- | --- | --- |
| GO:0033256 | I-kappaB/NF-kappaB complex | 4 | 5.53683 | 0.00055 | afc | J-Lat (Case) |
| GO:0005681 | spliceosomal complex | 112 | 1.24205 | 0.00074 | ks | J-Lat (Case) |
| GO:0030667 | secretory granule membrane | 59 | 0.566506 | 0.0032 | ks | J-Lat (Case) |
| GO:0022626 | cytosolic ribosome | 68 | 0.22662 | 0.00054 | ks | Jurkat (Control) |

**Table 2.**
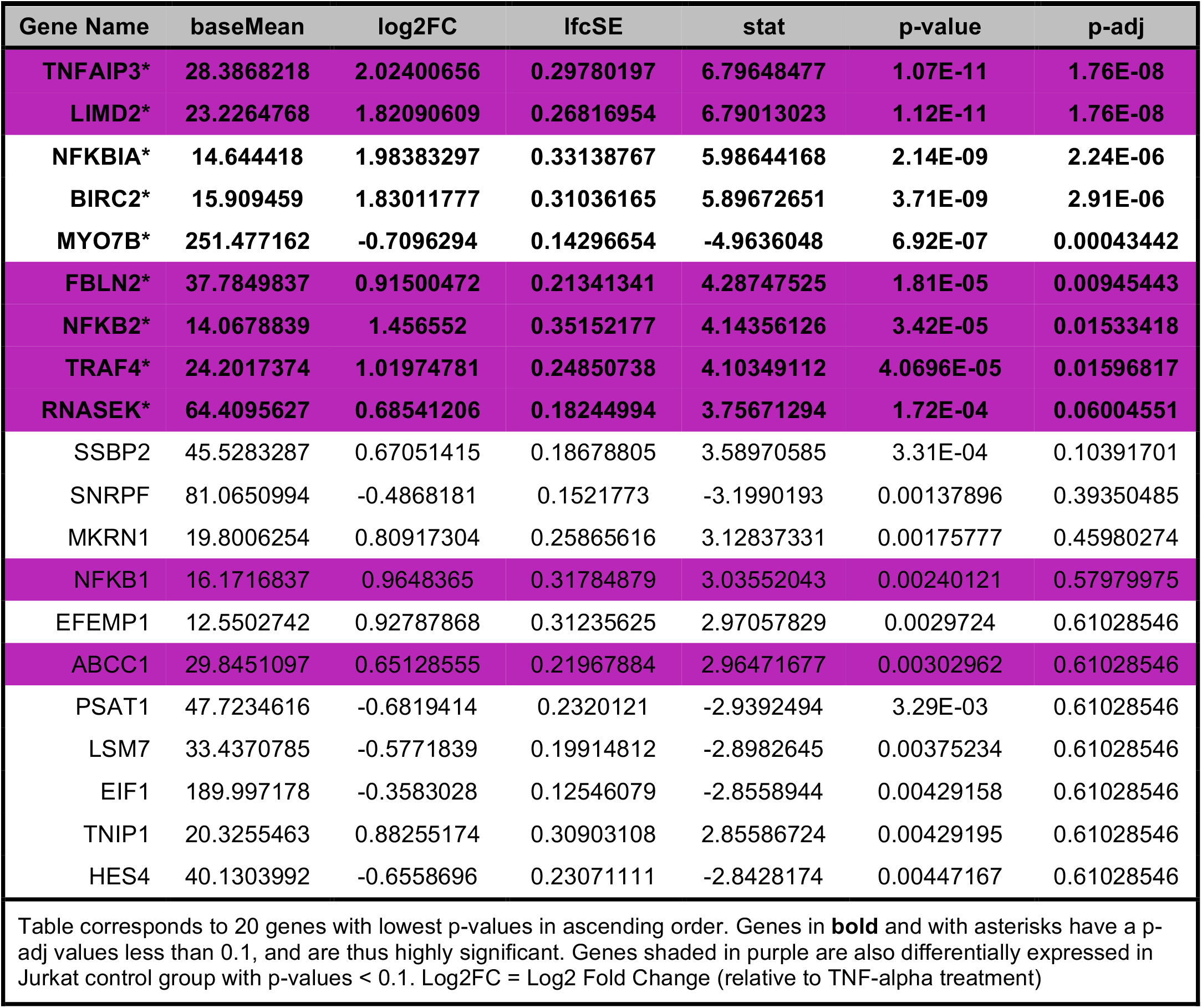
Differential Gene Expression in J-Lat 10.6 case group elicited by TNF-alpha.

To gain further insights into the specific transcript variants or isoforms eliciting gene expression changes, we plotted the TPM values of differentially expressed isoforms (DIE) with p-value<0.01 in the J-Lat case group **(Figure 5A)**. Hierarchical clustering shows two distinct populations, that are up- or down-regulated upon TNF-alpha induction. Those isoforms also found to be differentially expressed in Jurkat control group were highlighted in purple, and genes found to be highly significant (p-adj<0.1) are in bold and denoted with an asterisk. As consistent with the differential gene expression data, most of the highly significant DIE isoforms are upregulated upon TNF-alpha induction, with only a single isoform of PSAT1 being downregulated in this group. The DE isoform data confirms the involvement of NFKB-complex via significant 4-fold increases in relevant NFKBIA and BIRC2 isoform TPMs upon TNF-alpha treatment. Of particular note, is the highly significant (p-adj<0.1) paradoxical downregulation of a PSAT1 isoform given that previous studies have found this gene to be enriched during Tat-elicited cell proliferation in productive HIV infection (45) and during FOXO1-inhibition elicited latency reversal in HIV-infected CD4 T cells (46). This paradoxical result can be reconciled by close inspection of its exon connectivity **(Figure 5B)**, which reveals that the downregulated isoform (NM_021154.4) lacks Exon 8 and results in a variant with 6-to 7-fold lower activity compared to the standard NM_058179.4 isoform that retains this exon (47). The downregulation of the PSAT1 isoform lacking exon 8 that we observed in our data is consistent with the statistically-significant decrease in usage of the Exon 7 to 9 splice junction derived from our re-analysis of an independent and published Illumina dataset that used the J-Lat 10.6 clone and similar TNF-alpha induction conditions (**Supplementary Figure 11**) (31). Exon 8 contains a serine 331 residue which was shown to be phosphorylated by IKBKE, a known activator of NFKB pathway, and this modification results in a downstream activation of the serine biosynthetic pathway (SBP) to support cell proliferation (48). Besides providing a putative link between NFKB-complex and the SBP in a latency reversal context, the coupling of exon connectivity along with differential isoform expression shows the utility of full-length approaches to clarify seemingly paradoxical mechanisms of transcriptional regulation.

**Figure 5.**
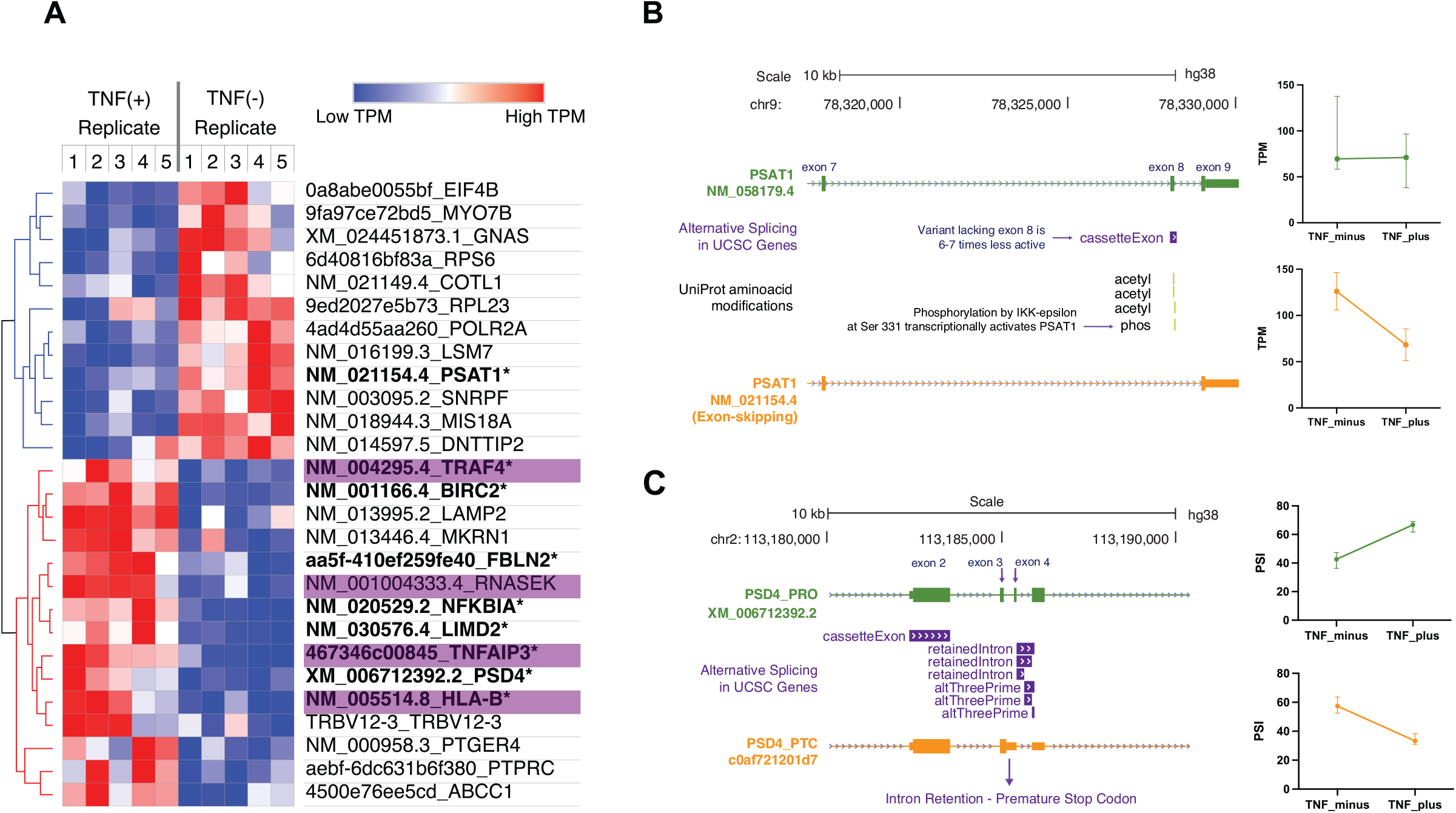
Differential Isoform Expression analysis shows putative HIV host factors PSAT1 and PSD4 are alternatively spliced in host cells upon HIV reactivation. **(A)** Heatmap showing hierarchically clustered TPM values for differentially expressed isoforms (p-value<0.1). Highly significant hits (p-adj<0.1) are in bold, while isoforms shaded in purple are also present in Jurkat control group. **(B)** PSAT1 isoform lacking functionally important exon 8 is differentially downregulated upon HIV reactivation with TNF-alpha. **(C)** Unproductive PSD4 isoform containing novel intron retention event is predominantly expressed in J-Lat cells prior to HIV induction. Upon HIV reactivation with TNF-alpha, intron retention event is downregulated and productive isoform is upregulated.

To further investigate changes in splicing as a response to TNF-alpha induced viral reactivation in host cells, we used the FLAIR DiffSplice module to call alternative splicing events from collapsed isoform clusters. An intron retention event between exon 3 and exon 4 in the PSD4 gene locus was found to be significantly (p-adj<0.05) modulated upon TNF-alpha induction in J-Lat 10.6 cells **(Figure 5C)**. This intron retention event, which is novel and not found in UCSC or SIB databases, was uniquely found in J-Lat 10.6 Case group and results in a premature termination codon which renders this transcript variant unproductive. DRIMSeq2 data was used to calculate the percent spliced in (PSI) of this intron retention event and showed the non-productive isoform was predominant in uninduced J-Lat (60% PSI value), but upon TNF-induction, the intron retention was downregulated resulting in 65% PSI in the productive isoform. This dynamic is concomitant with the robust induction (p-adj<0.1) of the productive XM006712392.2 PSD4 isoform upon TNF-alpha treatment as evidenced by its 2-fold increase in normalized isoform expression **(Figure 5A)**. PSD4 belongs to a family of Pleckstrin and Sec7 domain containing proteins (PSD or EFA6), which are associated with the plasma membrane (PM) and interact with ARF6 proteins via their Sec7 guanine exchange factor domain to regulate PM and endosomal traffic (49). ARF6 has been previously found to be a molecular determinant of HIV-1 Gag association with the PM (50) via its activation of PIP5K lipid modifying enzyme(51) which enhances PIP2 production, an acidic phospholipid which is specifically recognized by the highly basic region of HIV Matrix for anchoring into PM (52). Despite the wealth of evidence of an ARF6 interaction with Sec7 domain containing proteins, PSD4 has not been directly associated with productive HIV infection or evaluated for its regulation via an intron retention mechanism.

In addition to host cell transcriptional correlates, our approach also captures the HIV transcriptional signature that is concomitant to TNF-alpha induced viral reactivation in J-Lat 10.6 cells. Our Nanopore approach to quantify HIV transcripts performs favorably with competing datasets with regards to coverage evenness, with cross-comparison of spliced reads with an Illumina-sequenced TNF-alpha induced J-Lat 10.6 clone from an independent study showing uneven coverage across exons, and oversampling at splice junctions **(Supplementary Figure 12)**. Isoform clustering and collapse analysis across four Nanopore-sequenced replicates shows the capture of all canonical HIV splice sites and all multiexonic transcripts **(Figure 6A)**. These transcripts are divided into “Completely Spliced” (i.e. 2kb), and “Incompletely Spliced” (i.e. 4kb) classes based on the presence or lack of a D4-A7 splice event. However, unlike previous approaches (18,21), direct comparison of enrichment between any transcript is possible in our approach irrespective of transcript class **(Figure 6B)**. In addition to canonical HIV isoforms, our approach showed presence of a Nef isoform lacking canonical A5-D4 exon which despite retaining complete ORF, has not been previously observed. Additionally, a completely spliced variant of Vif was observed, which despite lacking the canonical intron retention between D4-A7, still contains a complete and undisrupted ORF upstream to this site. With regards to non-coding exons 2 and 3, these are present at much lower enrichment levels compared to previous studies (21), with non-coding exon 3 being more prevalent and associated with Rev/Nef/Tat/Env transcripts, and non-coding exon 2 being less prevalent and only associated with Tat and Nef transcripts. Gene assignment was based on a two-variables, with ORF proximity to the 5’ end of isoform being initial variable, followed by the presence of an undisrupted ORF. Using this system allows isoforms to be assigned to a gene unambiguously, particularly in cases of incompletely spliced transcripts containing A4 acceptors, where ORF proximity alone would impute an unproductive Rev isoform, instead of the likely productive Env/Vpu transcript. By classifying isoforms into likely expressed genes **(Figure 6C)** relative gene expression can be determined, with highest enriched genes being Nef, Rev and Env accounting for 45%, 27% and 20% of transcripts respectively. The high abundance of Nef and Rev, compared to the relatively low level of Tat is consistent with previous studies (21,53) and concordant with splice acceptor usage in our data **(Figure 6D)**. Moreover, the relatively high abundance of Rev is consistent with the requirement of this viral protein to oligomerize on RRE substrates to ensure the export of unspliced and partially unspliced transcripts out of the nucleus (54). As expected, the D1 splice donor shows highest usage followed closely by D4, the latter of which is consistent with the highest enrichment observed in transcripts containing the D4-A7 splice junction (i.e. fully spliced) **(Figure 6E)**. HIV splicing dynamics can be further explored with a splice junction matrix **(Figure 6F)**, showing all observed combinations of splice donor/acceptor junctions along with their enrichment, with D1-A5 and D4-A7 junctions being the most highly expressed junctions and correlating to Env to Rev/Nef transcripts respectively.

**Figure 6.**
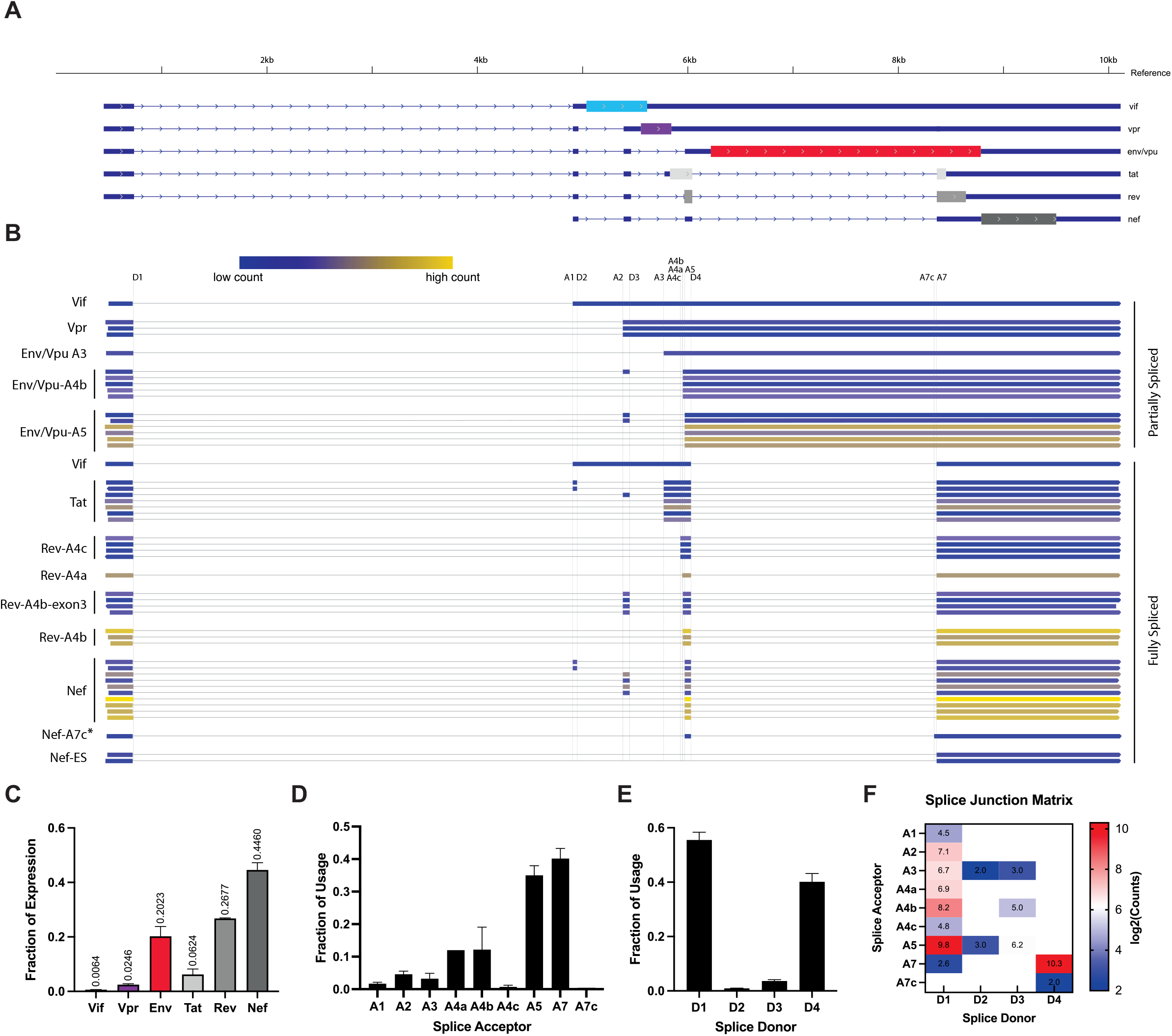
HIV transcriptional signature, gene expression and splice acceptor/donor usage for TNF-alpha induced viral reactivation in J-Lat 10.6 cells. **(A)** Idealized splicing structures of HIV genes and their CDS regions. **(B)** HIV multiexonic isoform clusters observed across four replicates are color coded based on count numbers, isoform clusters are annotated with likely gene expressed and differentiating splice acceptor junction. Non-canonical/novel isoforms are labeled with an asterisk **(C)** Gene expression fractions calculated based on counts obtained per isoform cluster, gene assignment based on proximity of ORF to 5’ end, and presence of undisrupted CDS. Splice **(D)** Acceptor and **(E)** Donor usage. **(F)** Splice junction matrix with log_2_ normalized counts shows association and frequency of specific splice donor/acceptor junctions.

## Discussion

In this study, we introduce and validate a full-length direct cDNA sequencing pipeline for the simultaneous profiling of poly-adenylated viral and host cell transcripts from unamplified cDNA. This approach is supported by the use of two high performing reverse transcriptases and Oligo-d(T) priming, coupled to a novel one-step chemical ablation of 3’ RNA ends, CASPR, which reduces rRNA reads and enriches poly-adenylated transcripts. We use this approach to simultaneously interrogate host and viral transcriptional dynamics within a full-length sequencing context in a relevant cell line model of HIV reactivation. This has allowed us to identify putative host factors of HIV transcriptional activation that contain exon skipping events (PSAT1) or novel intron retentions (PSD4). In addition, our full-length RNA-Seq pipeline is agnostic to sequencing methodology or library preparation approaches, and widely applicable for the study of viral transcription dynamics in host cells.

CASPR treatment in combination with MarathonRT were critical components in maximizing the quantitative capture of full-length host cell transcripts. The exact mechanism of CASPR-mediated improvements in obtaining full-length cDNA are beyond the scope of this manuscript; however, our data suggests that these improvements in priming specificity (via reduction of primer-independent products) are modulated by the 3’-OH ends of RNA inputs. The presence of non-specific cDNAs generated in a primer independent manner has been a largely overlooked artefact of reverse transcription. This has been cemented by the notion that exogenous DNA primers are an absolute requirement for reverse transcription, despite growing evidence of primer-independent cDNA generation in a variety of reverse transcriptases, which has been variously reported in the field as “false-priming”, “self-priming”, and “background priming” (55-58). Moreover, the fact that the CASPR reagent resulted in improvements in the performance of both MRT and SSIV despite their different origins, and in a variety of RNA inputs and priming modalities, points to RT initiation in absence of exogenous primer being a prevalent phenomenon. Primer-independent cDNA products are also a barrier in the study of replication dynamics of other RNA viruses where expression of negative strand intermediate transcripts is a hallmark of active viral replication, as is the case in Dengue Virus, West Nile Virus, Hepatitis C Virus, SARS-CoV2 and others (57,59-62). This suggests wide applicability of the CASPR reagent which, coupled with a suitable priming modality and a processive RT, could increase the breadth and sensitivity in the capture of full-length transcripts of interest in other relevant systems.

Given the polycistronic nature of HIV RNA, the full exon connectivity provided by this pipeline is a critical component in the unambiguous assignment of detected isoforms to a likely expressed gene or in the identification of novel splice variants. This is not a minor a problem for HIV, where a single intron retention event between two isoforms with seemingly identical splice junctions could result in expression of another viral gene. Full-length reads obtained in our pipeline allow straightforward isoform assignment and productivity analysis for the majority of HIV genes. However, the case of partially unspliced transcripts containing A4 or A5 splice sites constitutes an illustrative case where gene assignment can remain ambiguous even with full-length isoform information. Based on the premise that the closest ORF to the 5’ end of transcript constitutes the determinant factor in the gene that is expressed, partially unspliced isoforms containing A4 sites would translate to a unproductive Rev (since the CDS is disrupted by the D4/A7 intron retention), whereas those containing A5 would be translated as Env/Vpu. This ambiguity, however, is consistent with previous studies showing HIV co-opts the host cell translation machinery in non-canonical ways to further regulate its gene expression via leaky ribosomal scanning or ribosome shunting (63). Thus, ORF proximity to 5’ end is a necessary but not sufficient factor in determining which gene is eventually expressed from a particular splice variant. In these cases, we used the presence of a complete and non-disrupted CDS as a second prioritization scheme for gene assignment, whereby a partially spliced variant containing an A4 junction is likely to code for productive Env/Vpu and not an unproductive Rev (i.e., prioritization of longest ORF). Given the dynamic nature of HIV RNA secondary structure proximal to splice junctions(64) and its inhibitory role in ribosome scanning, future studies coupling splice variant detection with DMS-MaP secondary structure probing(34) might provide additional clarity on Rev and Env/Vpu translational regulation, while allowing additional variables for consideration of gene assignment and productivity analyses.

Despite the moderate sequencing depth used in our studies, the yield and coverage increases elicited by CASPR allowed sufficient capture of host cell transcript variants for biologically meaningful DGE/DIE analyses while also detecting all canonical splice junctions in HIV isoforms. Sequencing throughput in our studies was a function of the MinION sequencer used, which allowed for rapid method development and validation studies at the expense of number of reads (compared to some large scale transcriptomic studies of rare AS transcripts) (29). Any throughput constraints, can be easily addressed in future studies by adopting higher throughput platforms available from ONT, including the GridION and PromethION each with five- and 250-fold higher throughput. The higher sequencing depth provided by these platforms would enable detection of low-expression genes, resulting in higher sensitivity for rare genes, isoforms, or splice variants. An additional consideration in our platform hinges on the number of cells required for dispensing with PCR amplification, currently 50,000 cells are required to obtain sufficient total RNA. The required number of cells might not be unreasonable when using cultured cell lines, but when using primary cells or clinical samples, the requirement might be a limitation without further PCR amplification. For these types of samples, a cDNA amplification library preparation kit which attaches 5’ and 3’ adapters during RT can be used with CASPR-treated RNA inputs, followed by emulsion PCR with a single primer set and with a modest number of cycles to minimize PCR sampling bias (65), and allow for enrichment comparison between transcripts.

An interesting finding revealed by our study is the predominant intron retention event observed in the PSD4 locus of uninduced J-Lat cells, which results in expression of a truncated and inactive isoform due to a premature termination codon. The biological relevance of this AS event is not yet established; however, the role of other Sec7 domain containing proteins in targeting of viral components to the plasma membrane via its guanine exchange factor activity and interaction with ARF6 has been thoroughly documented (51). The reduction in expression of productive PSD4 could reduce the amount of active ARF6 and thus affect the balance of phosphatidylinositol that allows permissive assembly or entry of viral components proximal to the plasma membrane. However, intron retention events are widespread in cancer transcriptomes (66), and given the origin of J-Lat 10.6 cells from immortalized T cell leukemia PBMCs, the causal relationship between the modulation of PSD4 (and other AS isoforms) and HIV replicative capacity has to be thoroughly validated in primary cells.

In summary, we developed and systematically validated a full-length RNA-seq pipeline for assessing viral RNA transcript dynamics within a host cell transcriptome. This approach is supported by use of highly processive RTs, coupled with CASPR, a novel one-step CDS enrichment strategy that outperforms prevailing PolyA+ selection strategies in the breadth and sensitivity of capture of host cell and HIV transcripts. The simultaneous host and viral transcriptional signatures revealed in our approach can be used to interrogate transcriptional changes in response to a variety of viral induction methodologies, host gene manipulations (i.e. knockdown and knockouts), and viral sequence mutations, allowing greater granularity in the study of the interdependence of host and viral transcriptional regulation during infection with HIV and other RNA viruses.

## DATA AVAILABILITY

Sequencing data has been submitted to the NCBI Sequence Read Archive (SRA) under BioProject ID PRJNA801353

## Supplementary Figures and Tables

**Supplementary Figure 1.**
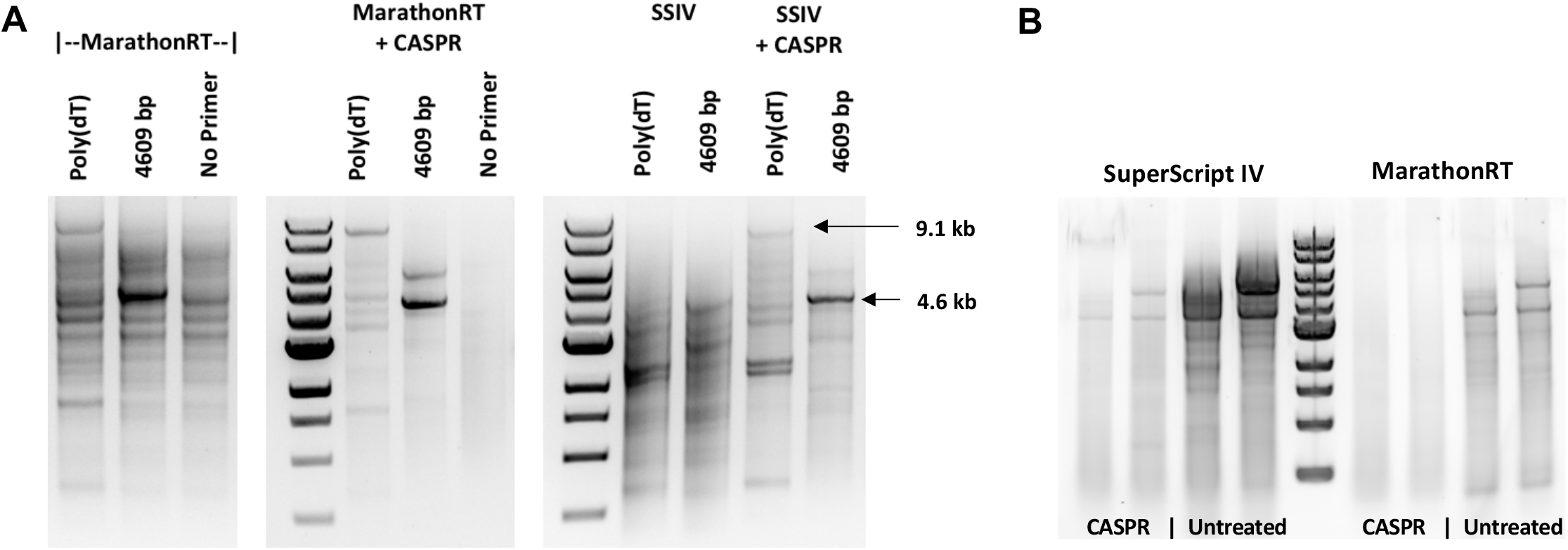
CASPR increases RT specificity of both MRT and SSIV when using a variety of RNA inputs and priming modalities. **(A)** Agarose gel electrophoresis of SSIV and MRT cDNA products from in vitro transcribed HIV-1 RNA inputs when using Oligo-d(T) or Gene-Specific Priming modalities **(B)** Agarose gel electrophoresis of SSIV and MRT cDNA products from HEK-293T total RNA when using Oligo-d(T) priming.

**Supplementary Figure 2.**
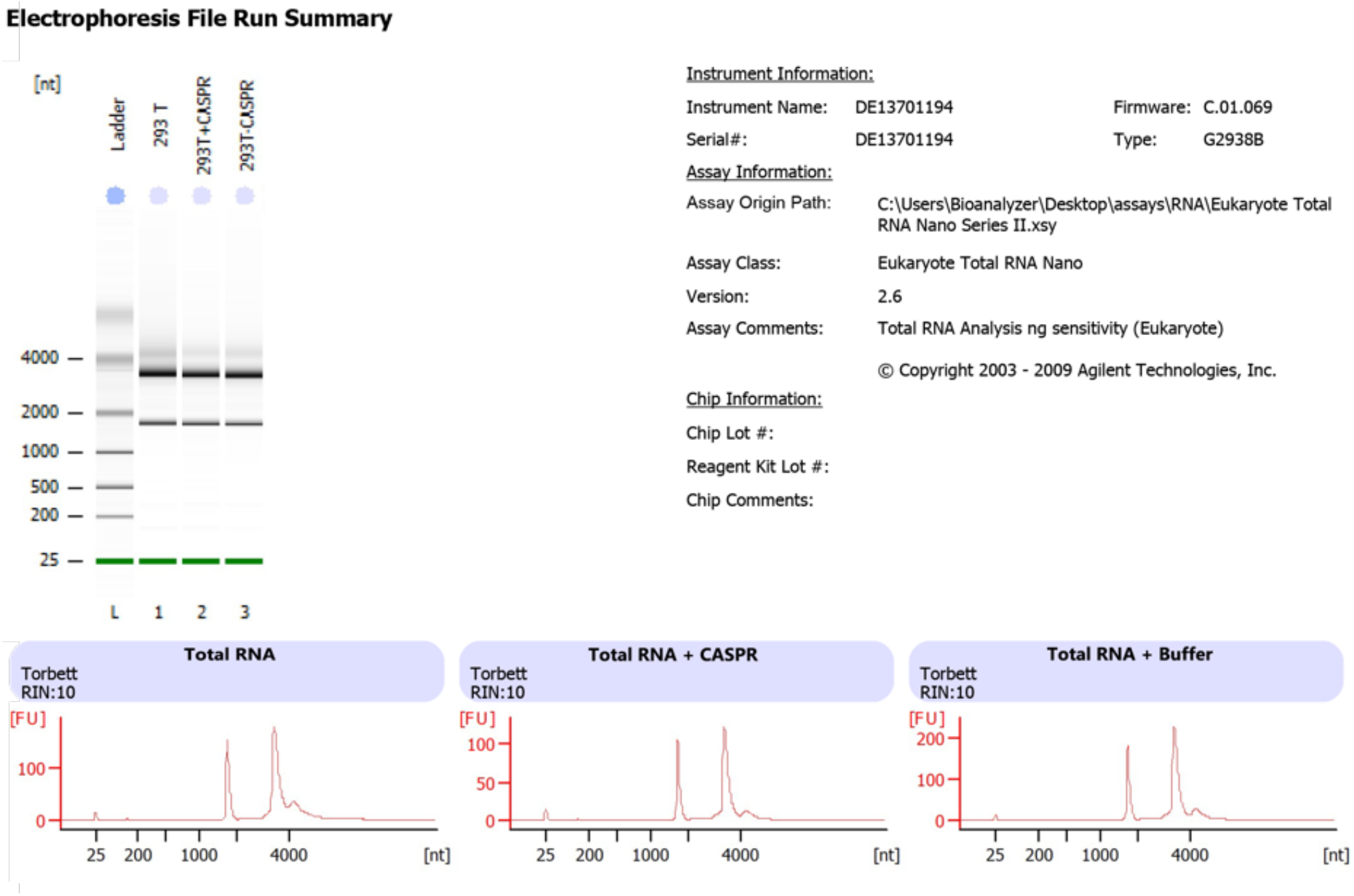
CASPR treatment does not compromise total RNA integrity as measured by Agilent RIN score. Run summary from Agilent Bioanalyzer Total RNA Nano Chip of untreated Total RNA, Total RNA with CASPR, and Total RNA with CASPR Buffer only. RNA Integrity Numbers (RIN) of 10 are reported for all samples, illustrating our RNA handling procedures and CASPR treatment, do not compromise the quality of RNA used in our pipeline.

**Supplementary Figure 3.**
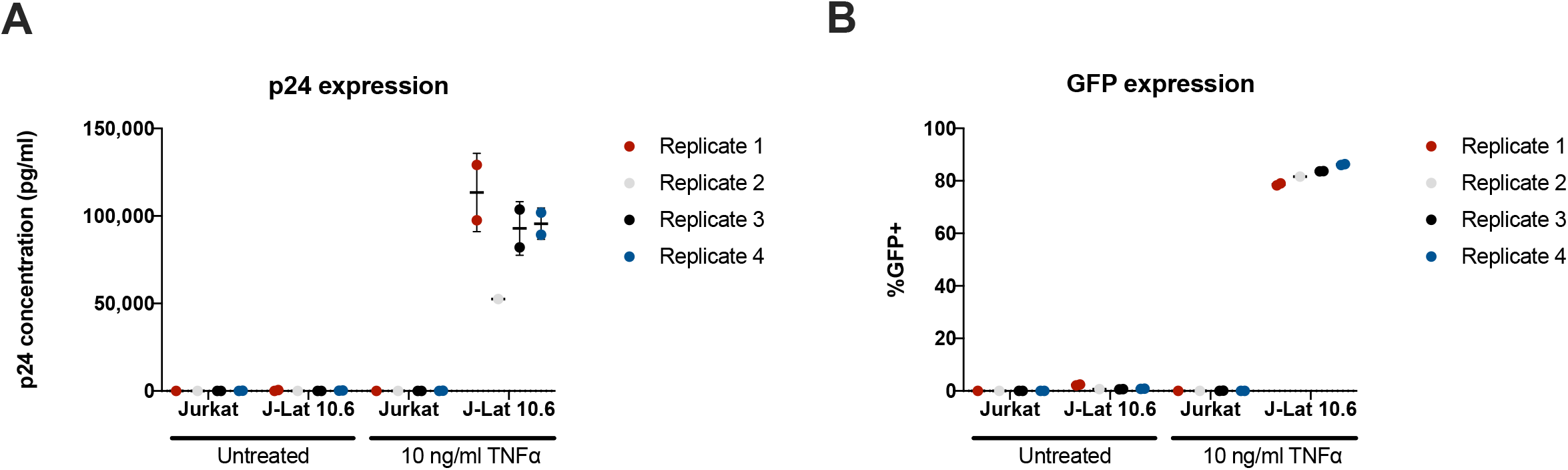
p24 and GFP% induction in J-Lat 10.6 cells after TNF-alpha induced latency reversal is nominal compared to previous studies. (A) p24 induction from viral supernatants of Jurkat and J-Lat cells activated with TNF-alpha. (B) GFP% induction for all samples, GFP% of 75-80% across J-Lat replicates

**Supplementary Figure 4.**
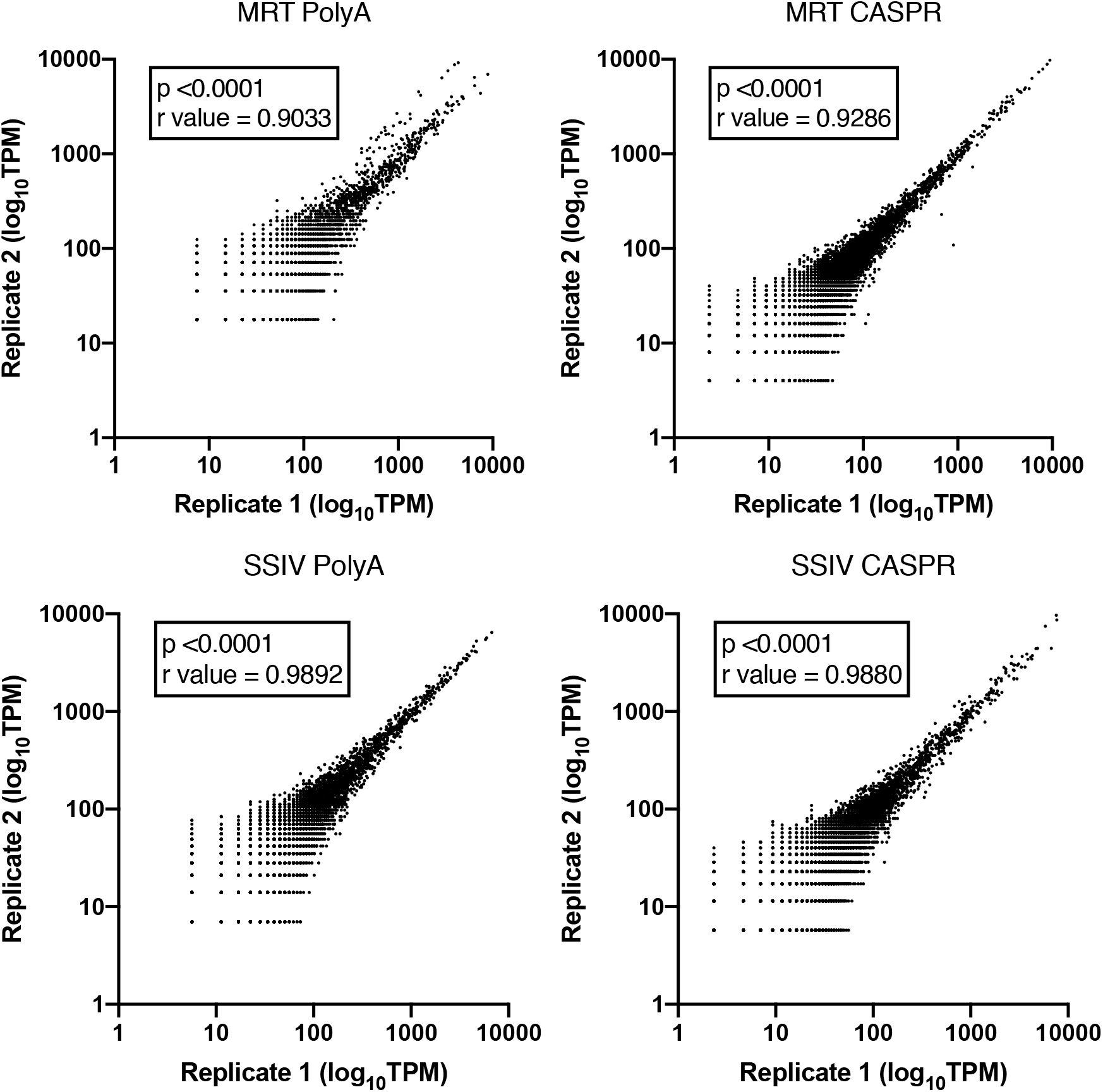
Reproducibility of PolyA and CASPR gene expression TPM values across replicates and treatments.

**Supplementary Figure 5.**
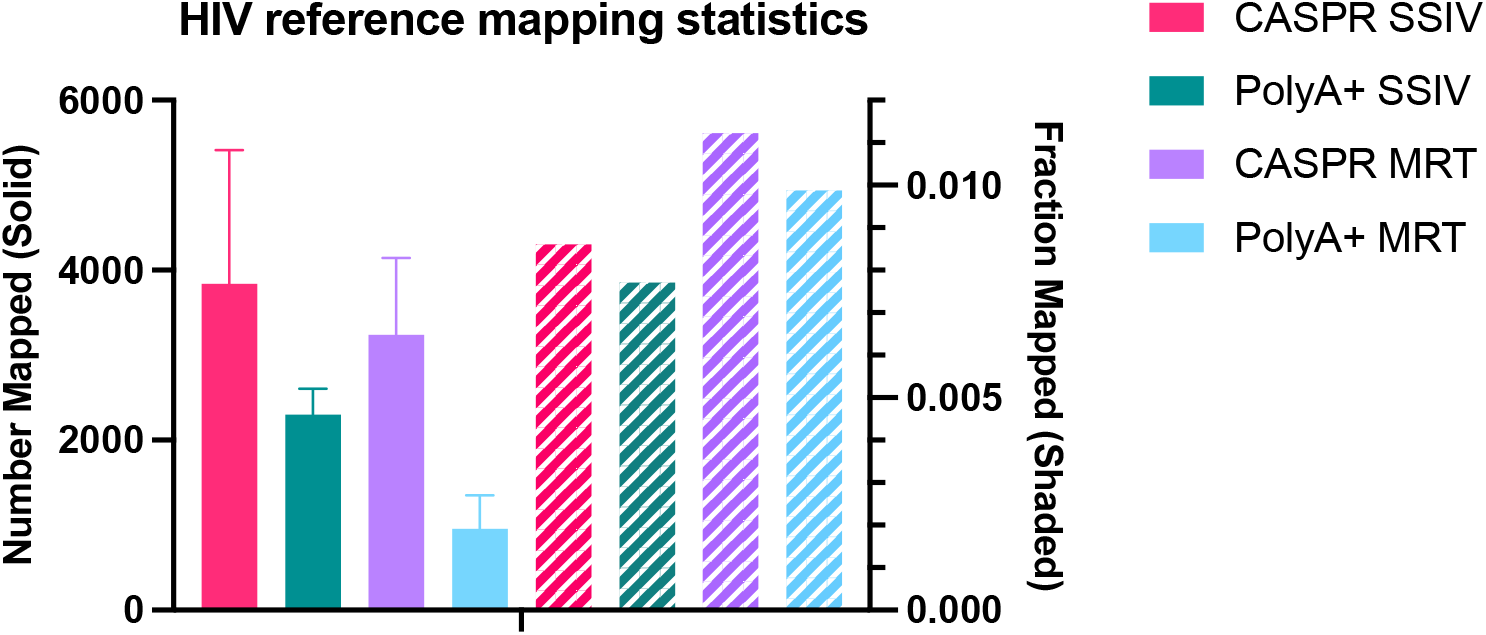
HIV mapping statistics. Number (solid) and fraction (shaded) of reads mapping to HIV reference sequence

**Supplementary Figure 6.**
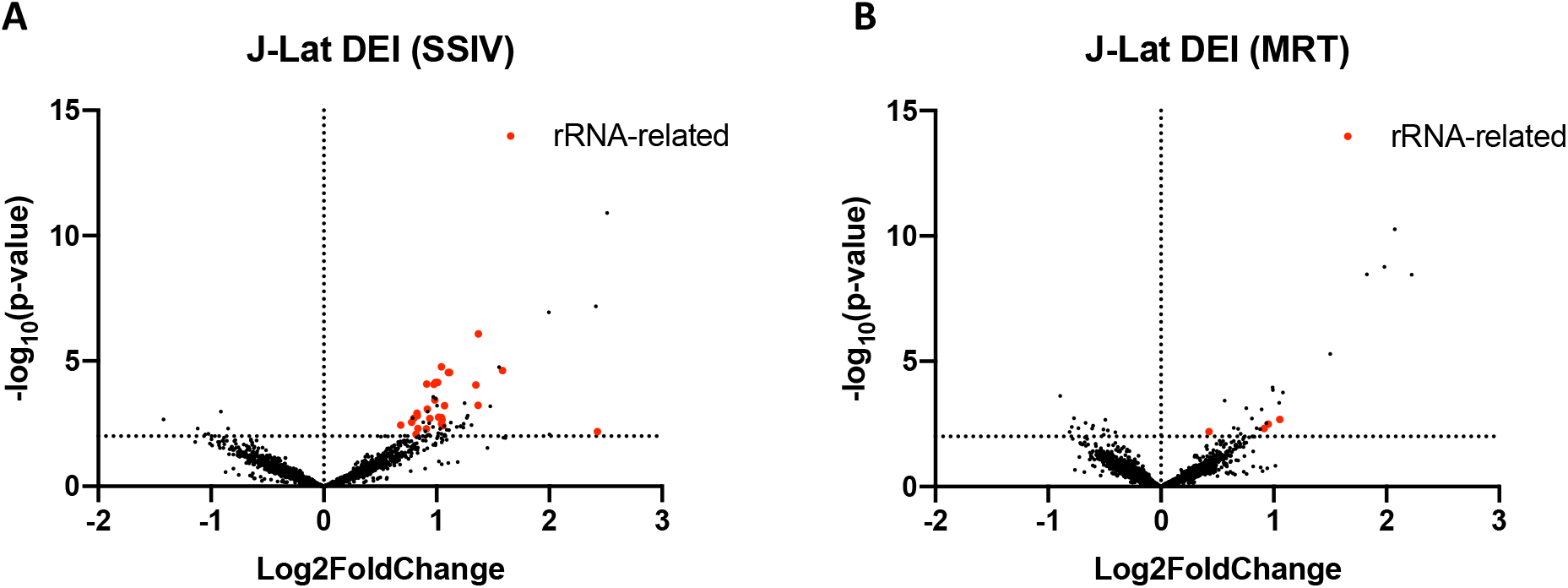
Pilot differential isoform expression (DIE) analysis using CASPR treated RNA inputs, SSIV shows high number of artefactual rRNA related hits compared to MRT. Volcano plots showing high number of rRNA-related DIE hits (red dots) when using (A) SuperScript IV compared to (B) MarathonRT

**Supplementary Figure 7.**
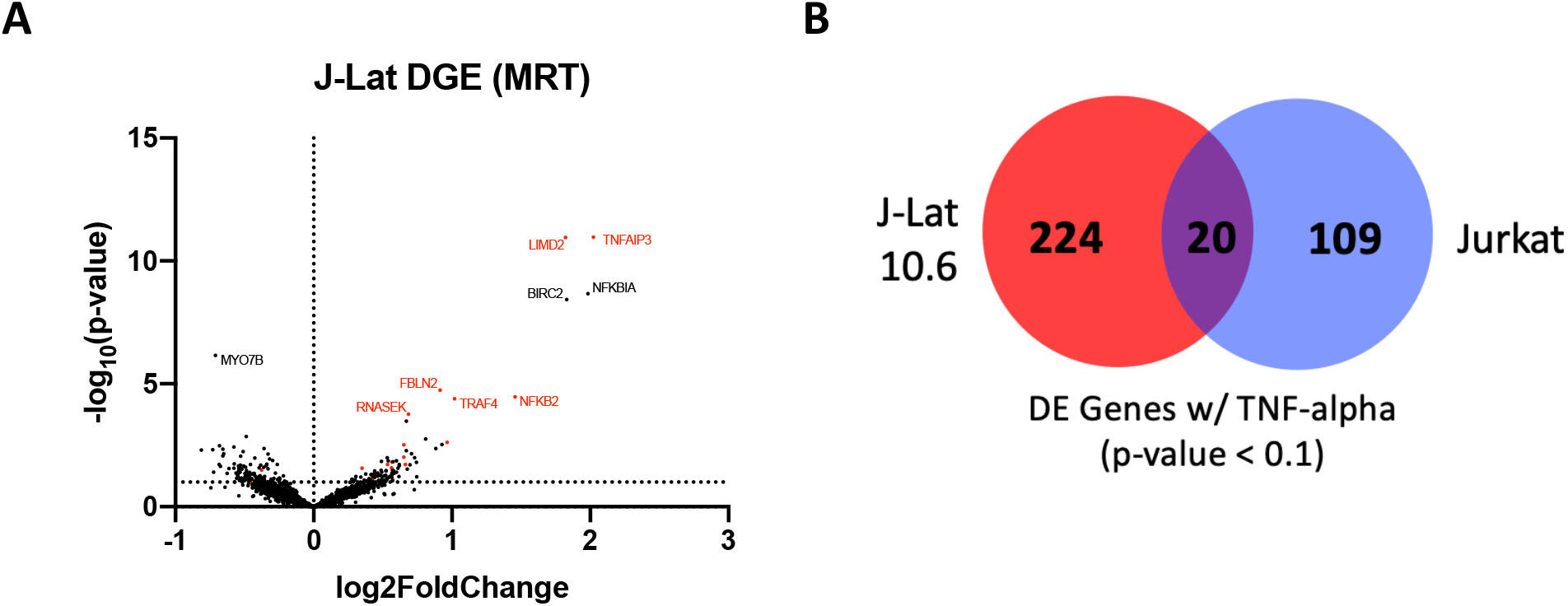
Overlap between genes differentially expressed upon TNF-alpha induction in J-Lat case group and Jurkat control Group. (A) Volcano plot showing differentially gene expression (DGE) hits with p-value<0.1, hits with gene names denote high significance (p-adj<0.1) and those in red denote genes present only in J-Lat case group (B) Venn diagram denoting number of genes showing DGE above threshold for case and control groups, along with degree of overlap.

**Supplementary Figure 8.**
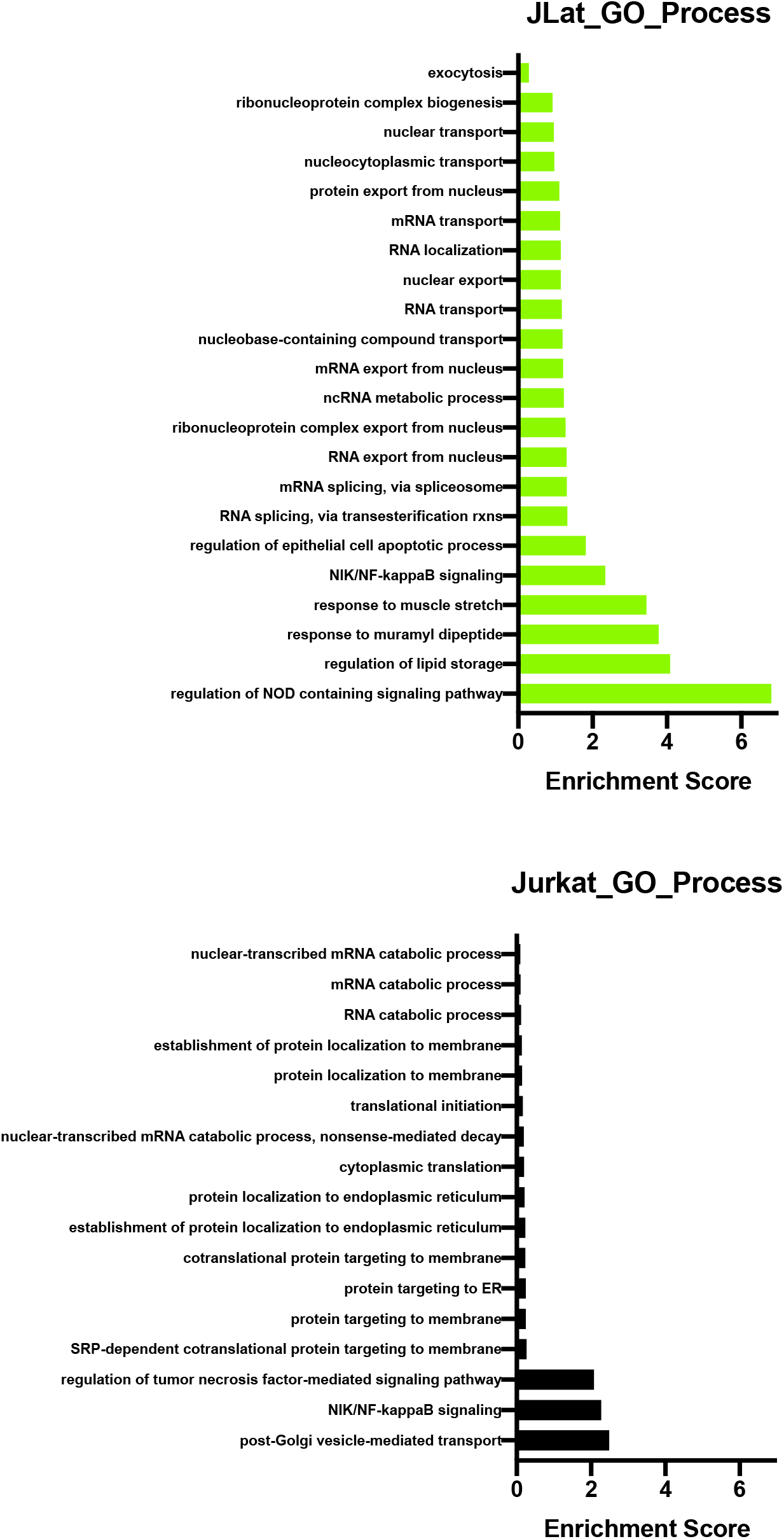
Pathway analysis (GO Biological Process) for Jurkat and J-Lat 10.6 based on log_2_Fold Change values TNF-alpha induction.

**Supplementary Figure 9.**
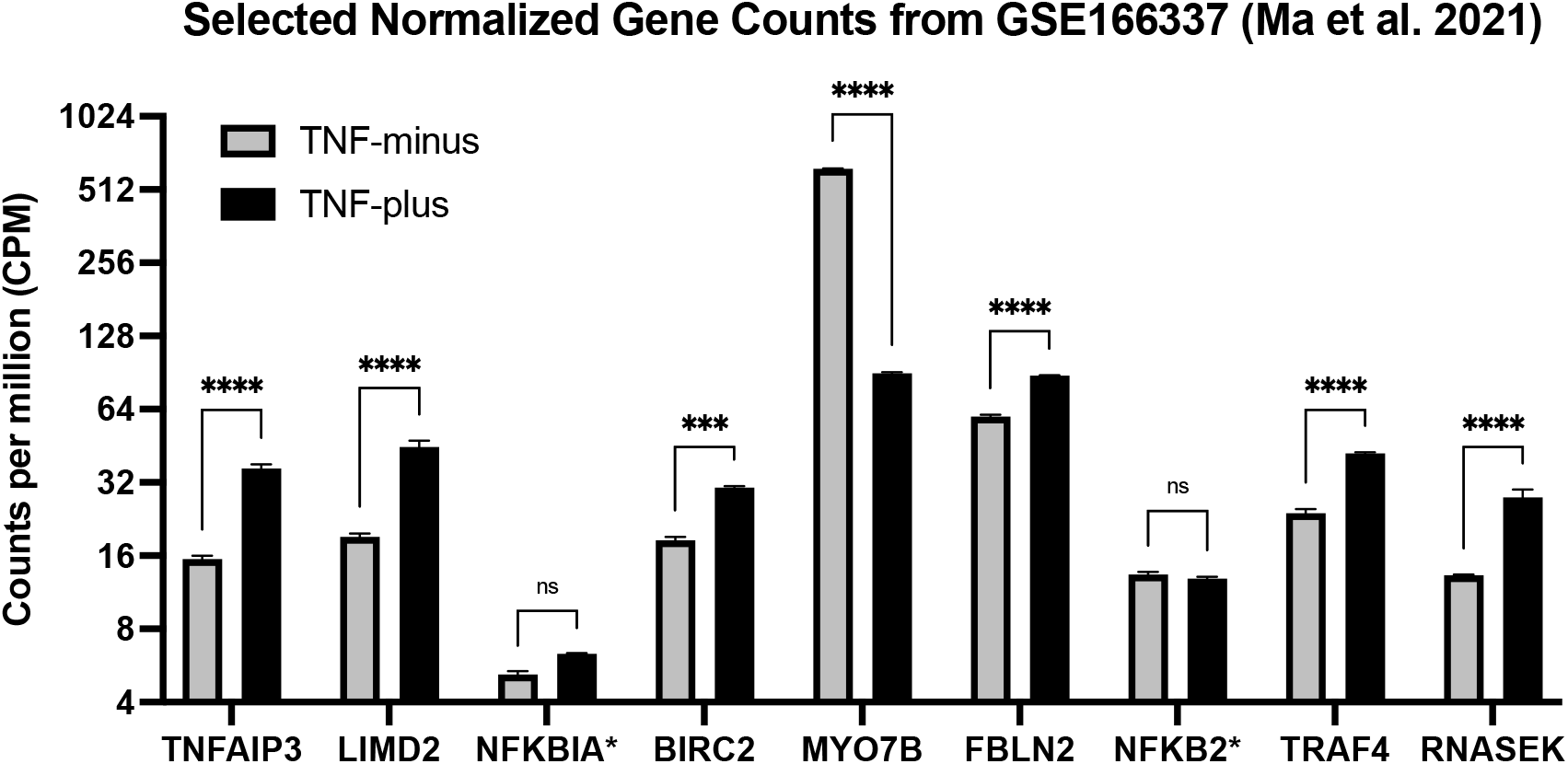
Selected normalized gene counts from Illumina-sequenced publicly-available data using same cell line clone and TNF-alpha induction conditions. There is high concordance with DGE hits from our data with p-adj<0.1. Genes with an asterisk denote genes not showing statistically-significant change in counts between TNF-plus and TNF-minus conditions. Re-analyzed data from Ma et al. 2021 (31).

**Supplementary Figure 10.**
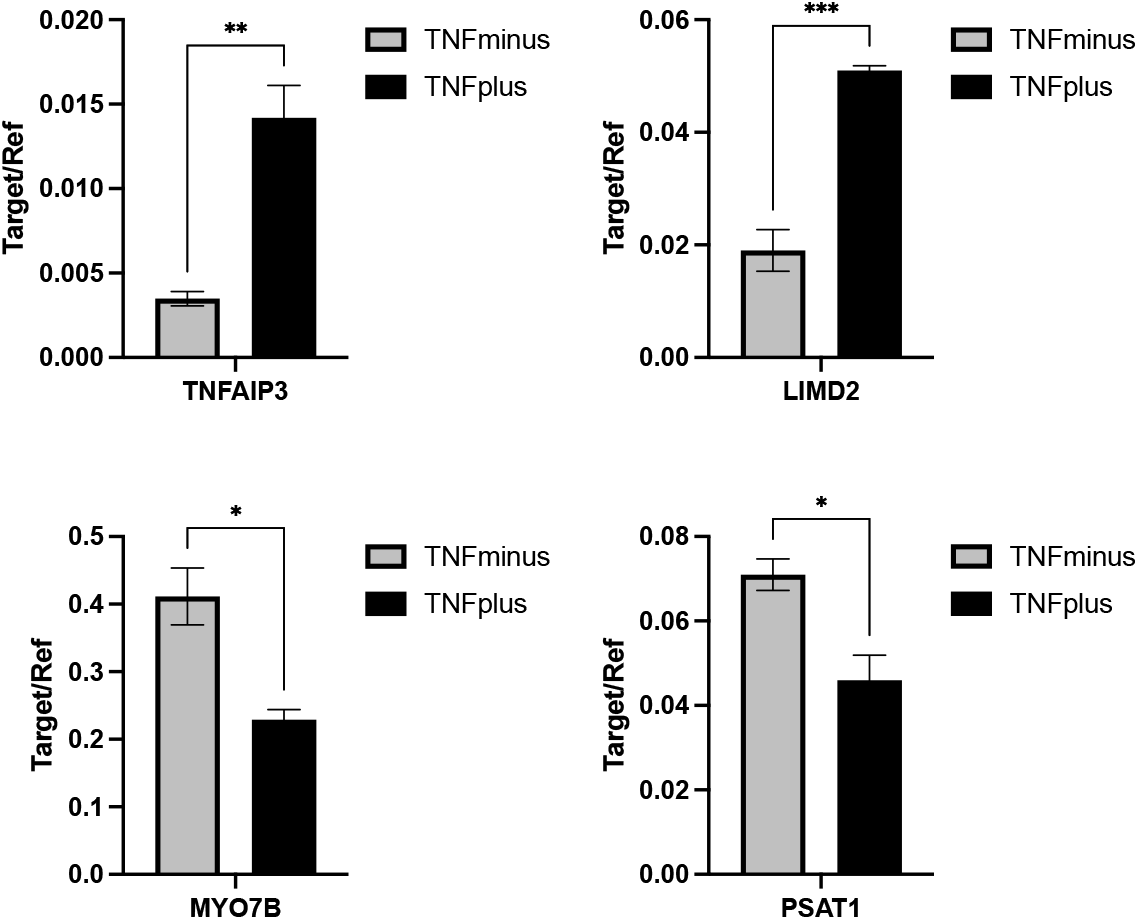
qPCR Validation of subset of significant DGE hits in TNF-plus and TNF-minus treatment groups. CASPR-treated RNA was reverse transcribed with MarathonRT, followed by second-strand synthesis. qPCR was carried out using PrimeTime qPCR Primer Assays (IDT) for the listed Targets and using Beta Actin (ACTB) as a Reference Gene (n=3 for TNF-plus, and n=4 for TNF-minus). Relative quantification results shown as ‘Target/Ref’, all values are means ± SEM. Statistical significance calculated with two-way ANOVA with Tukey multiple comparisons test, p<0.05(*), p<0.01(**), p<0.001(***).

**Supplementary Figure 11.**
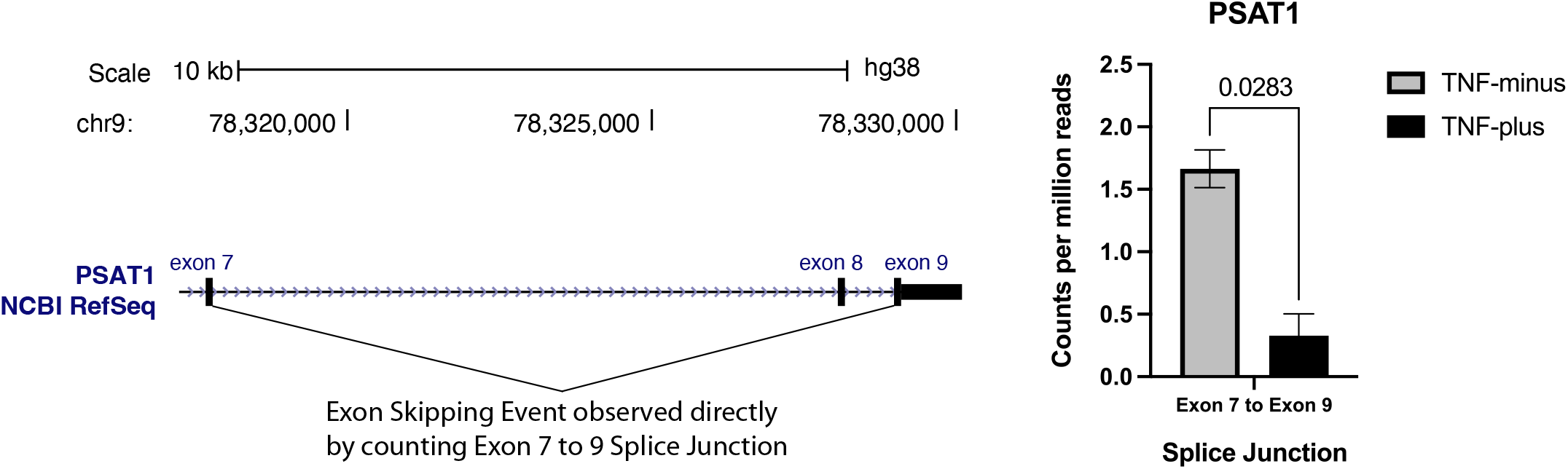
Exon 7 to Exon 9 Splice Junction Usage from PSAT1 gene from Illumina-sequenced publicly available that uses J-Lat 10.6 clone and TNF-alpha induction. Following TNF-alpha induction, there is a statistically significant (p<0.05) 3-fold reduction in CPM value of Exon 7 to 9 splice junction, which is highly concordant with the decrease of isoform lacking Exon 8 observed via ONT long-read sequencing. Re-analyzed data from Ma et al. 2021 (31)

**Supplementary Figure 12.**
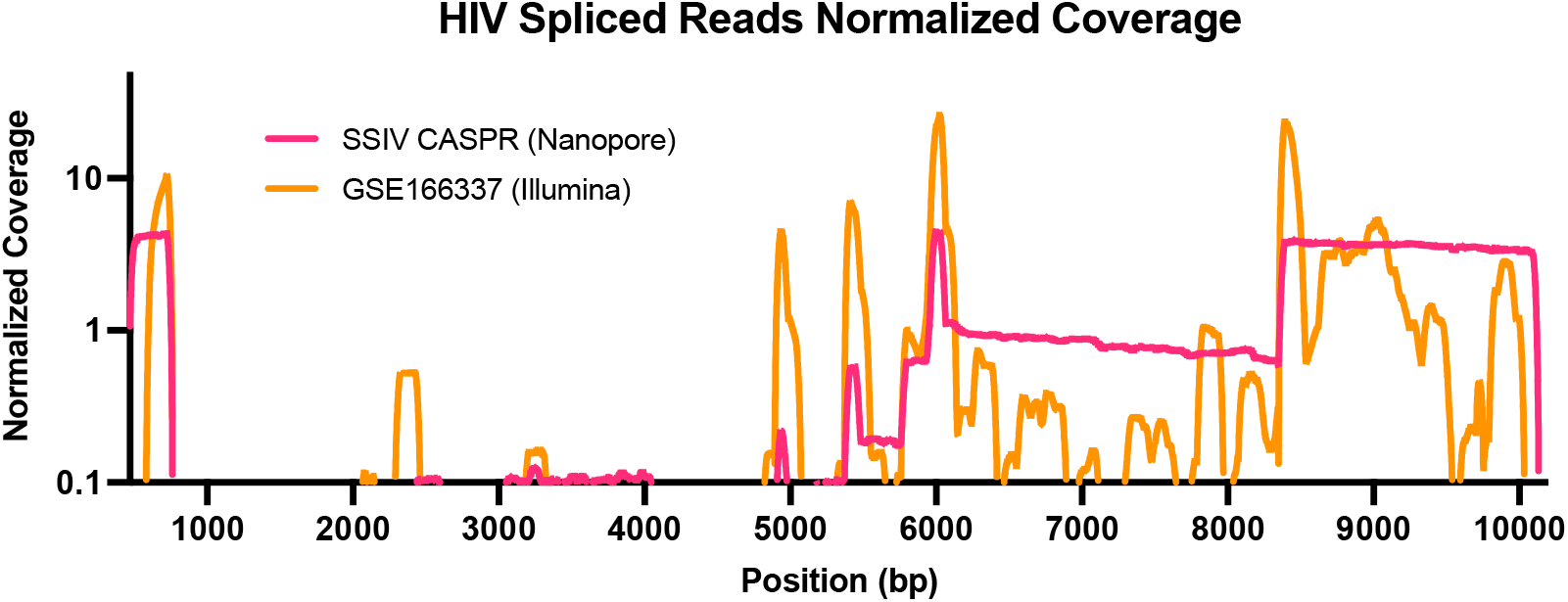
Normalized coverage comparison of spliced reads obtained with our long-read approach and an published Illumina dataset using TNF-alpha induced J-Lat 10.6 clone. To account for differences in read-depths between the datasets, coverage at each position was normalized to the average number of reads for the whole dataset, with coverage of lower or greater than 1 denoting under-and over-sampling respectively.

**Supplementary Table 1.** HIV Splice Junctions Captured.

| Splice ID | Position (bp) | Type | Notes |
| --- | --- | --- | --- |
| D1 | 743 | Conserved |  |
| D2 | 4962 | Conserved | Non-coding exon 2 |
| D3 | 5463 | Conserved | Non-coding exon 3 |
| D4 | 6045 | Conserved | vpr, tat, rev, nef, vif |
| A1 | 4913 | Conserved | Non-coding exon 2, vif |
| A2 | 5390 | Conserved | Non-coding exon 3, vpr |
| A3 | 5778 | Conserved | tat |
| A4c | 5937 | Conserved | Rev (acceptor C>T) |
| A4a | 5955 | Conserved | Rev (acceptor G>C) |
| A4b | 5961 | Conserved | Rev (major rev) |
| A5 | 5977 | Conserved | Nef, env/vpu (major Env) |
| A7c | 8346 | Novel |  |
| A7 | 8370 | Conserved | Vpr, tat, rev, nef |

**Supplementary Table 2.** Read Counts and QC metrics for J-Lat and Jurkat Sequencing Experiments.

| SAMPLE ID | RT<br>CONDITION | TNF<br>INDUCTION | CELL<br>LINE | READS | BASES | AVG.<br>LENGTH | MAX<br>LENGTH | N50 | PAIR ID* |
| --- | --- | --- | --- | --- | --- | --- | --- | --- | --- |
| 1212-SC-TP-JL | SSIV CASPR | TNFplus | J-Lat | 346,371 | 380,624,842 | 1,098.90 | 18,134 | 1,608 | 1 |
| 1212-MC-TP-JL | MRT CASPR | TNFplus | J-Lat | 321,743 | 308,475,236 | 958.8 | 23,212 | 1,479 | 1 |
| 1212-SC-TM-JL | SSIV CASPR | TNFminus | J-Lat | 413,354 | 470,128,670 | 1,137.40 | 21,715 | 1,689 | 2 |
| 1212-MC-TM-JL | MRT CASPR | TNFminus | J-Lat | 342,722 | 348,029,436 | 1,015.50 | 20,294 | 1,567 | 2 |
| 1211-SC-TP-JL | SSIV CASPR | TNFplus | J-Lat | 465,339 | 494,378,685 | 1,062.40 | 19,100 | 1,583 | 3 |
| 1211-MC-TP-JL | MRT CASPR | TNFplus | J-Lat | 302,825 | 271,089,416 | 895.2 | 23,629 | 1,400 | 3 |
| 1211-SC-TM-JL | SSIV CASPR | TNFminus | J-Lat | 512,040 | 549,127,704 | 1,072.40 | 18,445 | 1,565 | 4 |
| 1211-MC-TM-JL | MRT CASPR | TNFminus | J-Lat | 341,749 | 332,377,217 | 972.6 | 25,594 | 1,506 | 4 |
| 10232-SC-TP-JL | SSIV CASPR | TNFplus | J-Lat | 207,115 | 273,356,212 | 1,319.80 | 20,503 | 1,879 | 5 |
| 10232-MC-TP-JL | MRT CASPR | TNFplus | J-Lat | 297,355 | 289,320,116 | 973 | 31,113 | 1,344 | 5 |
| 10232-SC-TM-JL | SSIV CASPR | TNFminus | J-Lat | 315,294 | 291,268,332 | 923.8 | 28,699 | 1,307 | 6 |
| 10232-MC-TM-JL | MRT CASPR | TNFminus | J-Lat | 175,399 | 138,830,006 | 791.5 | 24,842 | 1,175 | 6 |
| 10231-SC-TP-JL | SSIV CASPR | TNFplus | J-Lat | 510,364 | 752,703,706 | 1,474.80 | 27,758 | 2,143 | 7 |
| 10231-MC-TP-JL | MRT CASPR | TNFplus | J-Lat | 516,248 | 536,827,969 | 1,039.90 | 27,422 | 1,486 | 7 |
| 10231-SC-TM-JL | SSIV CASPR | TNFminus | J-Lat | 282,062 | 302,421,954 | 1,072.20 | 23,163 | 1,596 | 8 |
| 10231-MC-TM-JL | MRT CASPR | TNFminus | J-Lat | 255,840 | 236,887,897 | 925.9 | 18,757 | 1,403 | 8 |
| 10232A-MC-TP-JL | MRT CASPR | TNFplus | J-Lat | 322,176 | 279,486,945 | 867.5 | 21,991 | 1,160 | N/A |
| 10232A-MC-TM-JL | MRT CASPR | TNFminus | J-Lat | 560,222 | 489,725,213 | 874.2 | 29,792 | 1,175 | N/A |
| 210-MC-TP-JU | MRT CASPR | TNFplus | Jurkat | 466,114 | 407,911,711 | 875.1 | 18,298 | 1,376 | N/A |
| 1211-MC-TP-JU | MRT CASPR | TNFplus | Jurkat | 318,408 | 307,642,349 | 966.2 | 17,699 | 1,475 | N/A |
| 1023-MC-TP-JU | MRT CASPR | TNFplus | Jurkat | 327,561 | 312,003,518 | 952.5 | 16,492 | 1,469 | N/A |
| 1023-MC-TM-JU | MRT CASPR | TNFminus | Jurkat | 274,758 | 242,287,201 | 881.8 | 21,416 | 1,357 | N/A |
| 210-MC-TM-JU | MRT CASPR | TNFminus | Jurkat | 415,924 | 382,228,854 | 919 | 15,636 | 1,435 | N/A |
| 1211-MC-TM-JU | MRT CASPR | TNFminus | Jurkat | 218,935 | 215,360,547 | 983.7 | 16,965 | 1,511 | N/A |

\* Samples sharing the same Pair ID originate from the same biological specimen

